# Experimental enhancement of structural heterogeneity in forest landscapes promotes multidimensional hoverfly diversity

**DOI:** 10.1101/2025.08.22.670311

**Authors:** Clàudia Massó Estaje, Julia Rothacher, Ante Vujić, Marija Miličić, Anne Chao, Oliver Mitesser, Jörg Müller, Alice Claßen, Ingolf Steffan-Dewenter

## Abstract

1. The homogenization of temperate forests due to intensive management has led to biodiversity loss at local and landscape scales, threatening species persistence and ecosystem functions. However, experimental evidence on how structural heterogeneity among forest patches influences diversity at the landscape scale is lacking. Here, we test whether enhancing structural heterogeneity, via diverse deadwood enrichment and canopy gap creation treatments (Enhancement of Structural Beta Complexity, ESBC), can increase hoverfly diversity within landscapes, and elucidate the contributions of local (α-) diversity enrichment and turnover (β-diversity) to this effect.
2. We conducted a large-scale forest experiment in 11 regions across Germany. Each region included two districts, representing small forest landscapes. In one district, we implemented patches with ESBC treatments, while in the other, serving as the control, we established patches without ESBC. Hoverflies were sampled in three seasonal intervals and across, in total, 234 forest patches (50 x 50 m) using pan traps. We applied a new integrative meta-analytic framework that incorporates sample completeness to quantify taxonomic, functional, and phylogenetic diversity (TD, FD, PD) using Hill numbers at α, β, and γ scales.
3. All three γ-diversity dimensions - TD, FD, and PD - were significantly higher in structurally heterogeneous forest landscapes than in homogeneous ones. The strongest effects were observed for TD, indicating functional and phylogenetic redundancy among species. Effect sizes declined with increasing order of Hill numbers, suggesting that rare species benefit most from structural heterogeneity.
4. In most regions, γ-diversity gains were driven by increases in α-diversity rather than β-diversity, highlighting the importance of interventions to increase local structural complexity. However, several regions also showed elevated β-diversity, indicating context-dependent effects of spatial heterogeneity.
5. Our results provide the first experimental evidence that enhancing structural heterogeneity at the landscape scale can restore multi-dimensional insect diversity in temperate forests. They underscore the management value of ESBC as a scalable tool to restore biodiversity, increase ecological resilience, and counteract biotic homogenization in production forests.

## Introduction

Global biodiversity is declining at an alarming rate, with forest ecosystems being particularly vulnerable to human-induced pressures such as intensive management (McGill et al., 2015). In Central Europe, timber-focused forestry has structurally homogenized temperate forests by reducing canopy heterogeneity and deadwood availability (Eckelt et al., 2018; Ódor et al., 2006). This simplification contributes to biodiversity loss at both local (α) and landscape (γ) scales and drives biotic homogenization, the reduction in community dissimilarity (β-diversity), which can exceed the loss of species richness alone (McGill et al., 2015). Such patterns reduce ecosystem resilience and underscore the need for management strategies that promote structural heterogeneity across spatial scales (Gonçalves-Souza et al., 2025).

In agricultural landscapes, studies have shown that increasing structural heterogeneity, e.g. through measures such as diverse crop types, field margins, or hedgerows, can substantially enhance biodiversity by providing a variety of habitats and resources (Martin et al., 2019; Redlich et al., 2018). These insights have inspired research demonstrating that landscape-scale heterogeneity can maintain species richness and β-diversity across different land-use contexts (Tscharntke et al., 2012). In forests, structural heterogeneity among patches also plays a central role in biodiversity maintenance by creating a mosaic of microhabitats with diverse microclimates and resources (Thomsen et al., 2022). This includes among-patch variability in canopy cover, vertical layering, and deadwood availability (Uhl et al., 2024).

While effects of local, i.e. within-patch, structural simplification on α-diversity are well documented (Barnes et al., 2016; Mori et al., 2018), it remains poorly understood how much γ-diversity is shaped by α-versus β-diversity, particularly in the even-aged homogeneous stands that dominate temperate forests (Hilmers et al., 2018; Uhl et al., 2024). Earlier work in Central European forests has also shown that tree species diversity can drive strong α-, β-, and γ-diversity responses of arthropods, underscoring the role of patch-level compositional turnover for regional diversity (Sobek et al., 2009). In contrast to agricultural systems, experimental studies testing the biodiversity effects of spatial heterogeneity at the forest landscape remain scarce.

Recent forest management approaches aim to address these shortcomings through the Enhancement of Structural Complexity (ESC), which increases within-stand heterogeneity by retaining deadwood or opening the canopy (Heidrich et al., 2020; Schuldt et al., 2019). However, landscape-scale species loss cannot be addressed by local interventions alone. Building up ESC, the Enhancement of Structural Beta Complexity (ESBC) approach promotes β-diversity by increasing variation in structural attributes across forest patches (Müller et al., 2023). This landscape-scale heterogenization can foster different community assemblages through ecological mechanisms such as species sorting, habitat complementarity, and mass effects (Oehri et al., 2017; Perović et al., 2015). Additionally, natural disturbances such as windthrow, drought or insect outbreaks offer opportunities to reintroduce spatial heterogeneity and associated biodiversity benefits.

Species richness remains a widely used indicator to investigate how structural heterogeneity affects species communities; however, it overlooks functional and phylogenetic components of diversity, and without sample size standardisation, comparability among different habitat types can be biased (Chao & Jost, 2012). This study uses incidence data, for which Hill numbers offer a major advantage (Hill, 1973). Hill number is parameterized by an order q, which determines sensitivity to species frequency and provides a continuum of diversity measures that distinguish the influence of rare (q = 0), moderately common (q = 1), and dominant (q = 2) species (Chao et al., 2014; Colwell et al., 2012). Further, biodiversity assessments that ignore sampling completeness can lead to biased comparisons, as treatments may differ in how much of the regional species pool they capture (Kortmann et al., 2025). Methods based on rarefaction and extrapolation (iNEXT framework) allow adjustment for sample coverage (Chao et al., 2014). Combined with Hill numbers, this approach permits standardised, ecologically meaningful comparisons of diversity responses across treatments and biodiversity dimensions.

We applied the iNEXT framework to assess taxonomic (TD), functional (FD), and phylogenetic diversity (PD) (Chao et al., 2021), which provide complementary insights into how community composition reflects ecological roles and evolutionary history. While functional diversity reflects differences in species traits, phylogenetic diversity accounts for shared ancestry, which can capture unmeasured traits or evolutionary potential (Faith, 2016).

Hoverflies (Diptera: Syrphidae) were chosen as a model group for their ecological importance as pollinators and biological control agents, their sensitivity to microhabitat structure, and their strong responses to variations in canopy cover, microclimate and resource availability (Moquet et al., 2018; Popov et al., 2017). Their diverse life-history traits and phylogenetic breadth make them ideal for testing responses across TD, FD and PD.

We conducted a large-scale experiment in 11 forest regions across Germany, using a paired design comparing homogeneous (control) and heterogeneous (ESBC-enhanced) landscapes. Through manipulations of canopy cover and deadwood, we simulated natural successional dynamics to test how increased structural heterogeneity affects multiple biodiversity dimensions. We developed a meta-analytic framework that allowed direct comparisons across TD, FD and PD, and partitioned γ-diversity into its α and β components.

We hypothesize: (1) that ESBC interventions increasing forest landscape heterogeneity will enhance γ-diversity in TD, FD and PD compared to production forests; (2) that responses will be strongest in TD due to functional and phylogenetic redundancy; (3) that rare species will particularly benefit, leading to stronger effects at q = 0 than at q = 1 or 2; and (4) we expected that β-diversity will be a more important driver of γ-diversity than α-diversity due to the spatial structure of the interventions.

## Methods

### Study design

This study was conducted within the BETA-FOR experimental framework (Müller et al., 2023), spanning 11 temperate broadleaf production forest across six German regions: Lübeck (L11), Saarland (S10), Hunsrück (H09), Passau (P08), the Bavarian Forest National Park (B04-B07) and Sailershausen (U01-U03) (Fig. 1). Each forest was divided into two 10-20 ha districts, with 9 or 15 patches (50 x 50 m) established per district, totalling 234 patches.

**Figure 1.**
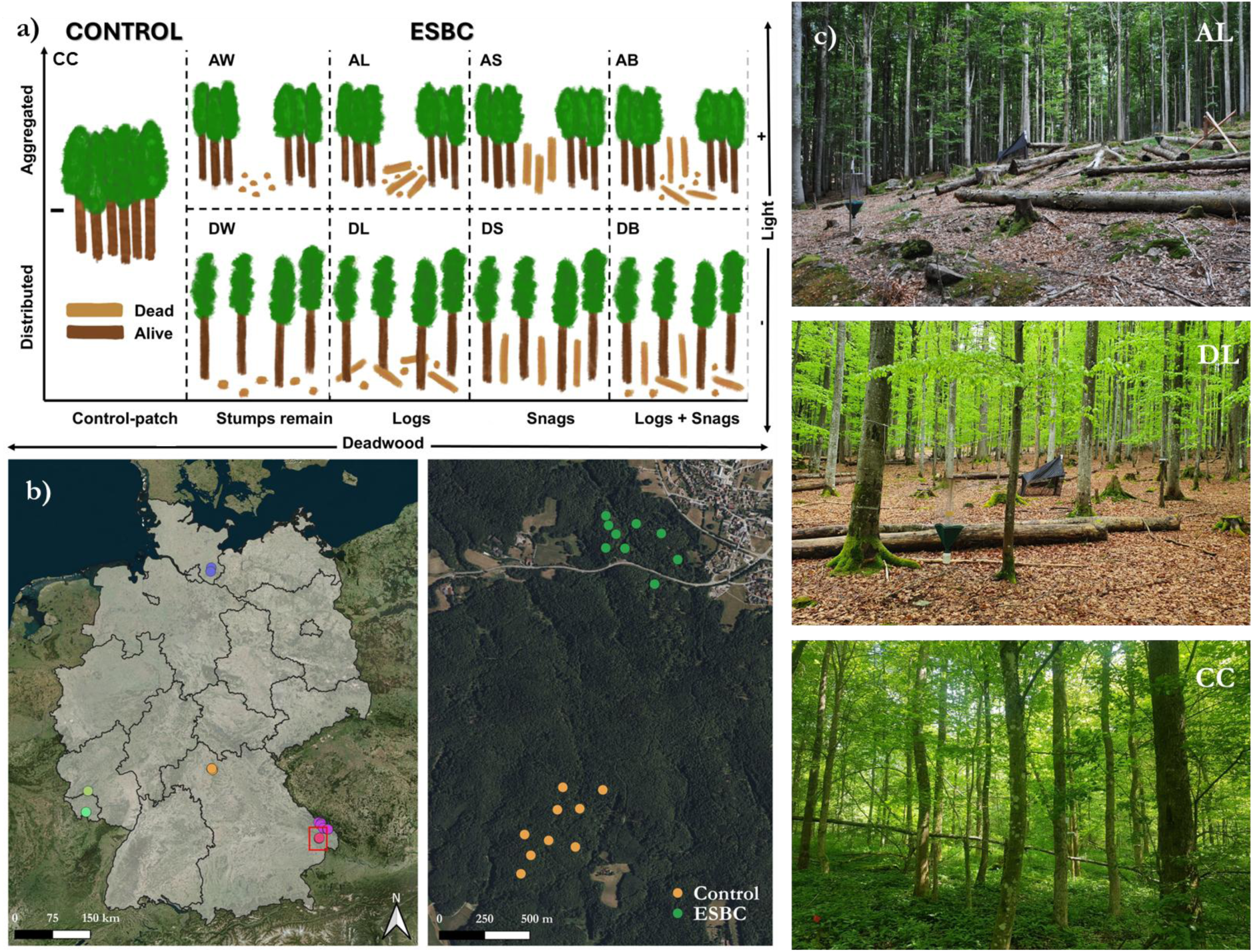
Study design of the BETA-FOR experiment: Treatments to Enhance Structural Beta Complexity (ESBC) manipulated forest structure via light and deadwood variation. We show 8 of the 14 experimental treatments (see Table 1); the remaining were only applied at the intensive study site (Sailershausen). Treatments vary along two axes: aggregation - Aggregated (central 30 m) or Distributed (across entire patch); and deadwood type - W: stumps; L: logs + stumps; S: snags; B: all three (stumps, logs, snags). (b) Map of forest districts showing Control and ESBC treatment locations. (c) Example photos of treatments across regions.

**Table 1.** Experimental treatments applied in the ESBC districts. Treatments were either aggregated (structural modifications in the central 30 m) or distributed (modifications across the entire patch). Interventions involved varying levels of tree removal and retention: total removal (R), retention of stumps (W), crowns (K), logs + stumps (L), snags (S), all woody structures (B), or creation of habitat trees (H). Some treatments were exclusive to Sailershausen.

| **Treatment** | **Aggregation Type** | **Description** | **Regions** |
| --- | --- | --- | --- |
| **AR** | Aggregated | Total tree removal | Sailershausen |
| **DR** | Distributed | Total tree removal | Sailershausen |
| **AW** | Aggregated | Tree removal, stumps remain | All regions |
| **DW** | Distributed | Tree removal, stumps remain | All regions |
| **AK** | Aggregated | Trees removed, crowns remain | Sailershausen |
| **DK** | Distributed | Trees removed, crowns remain | Sailershausen |
| **AL** | Aggregated | Logs and stumps remain | All regions |
| **DL** | Distributed | Logs and stumps remain | All regions |
| **AS** | Aggregated | Snags remain | All regions |
| **DS** | Distributed | Snags remain | All regions |
| **AB** | Aggregated | Stumps, logs, and snags remain | All regions |
| **DB** | Distributed | Stumps, logs, and snags remain | All regions |
| **AH** | Aggregated | Habitat trees created | Sailershausen |
| **DH** | Distributed | Habitat trees created | Sailershausen |

In each forest pair, one district – the ESBC district – underwent structural interventions to enhance structural β-complexity. Treatments altered canopy cover and deadwood features to mimic natural forest succession (Müller et al., 2023), affecting ∼30% of basal area per patch in varied ways to create heterogeneity. Table 1 summarizes the applied patch-level treatments.

The paired control district received no additional interventions, maintaining homogeneous management. Forests differed in soil, climate, and ownership, representing a range of temperate forest types from acidic beech-spruce-fir to mixed beech-oak stands.

### Hoverfly sampling

Hoverflies were sampled in all 234 BETA-FOR patches using standardized pan trapping. Each patch had three clusters of UV-reflective yellow, white and blue pan traps, placed along a transect with at least 15 m between clusters. Traps were filled with 400 mL of water and unscented detergent, left in the field for 48 hours, then emptied and preserved in 85% ethanol. All specimens were sorted and identified to species.

Sampling occurred in April, June, and July over two years. In 2022, 180 patches were sampled across the University Forests (U01–U03) and Bavarian Forest (B04–B07, P08). In 2023, 54 additional patches were sampled in Saarland, Hunsrück, and Lübeck (H09, S10, L11). Each patch was sampled with three traps across three periods, giving species incidence frequencies from 0 to 9.

### Trait and sequence data

Trait data for hoverfly species were compiled from different sources, including published data (Speight et al., 2021), expert opinion-based data, and fieldwork experience, spanning 35 years regarding biological and ecological characteristics of species across Southeast Europe. Eleven ecologically relevant traits were selected, including larval microhabitat and food type, larval development duration, number of generations per year, larval inundation tolerance, flight period, geographic distribution, adult body size, flying ability, adult microhabitat, and tolerance to human impact (Table S4).

For phylogenetic diversity (PD) analysis, COI-5P gene sequences were mainly retrieved from the Barcode of Life Data System (BOLD) using the bold_seq function in the bold R package (Dubois & Chamberlain, 2025). A phylogenetic tree was constructed using maximum likelihood via the TreeLine function in the DECIPHER R package (Wright, 2016), which models gene evolution by point mutations to estimate topology and branch lengths. The tree’s accuracy was validated against published phylogenies to ensure reliability for PD calculations.

### Calculation of TD, FD, PD

All analyses were conducted in R (v4.3.1; R Core Team, 2023). We compared TD, FD and PD between ESBC and Control districts across sites and diversity orders (Hill numbers and their generalizations to PD and FD, q = 0, 1, 2).

Effect sizes were calculated as the difference in diversity between ESBC and Control districts, with positive values indicating a higher diversity in ESBC patches. To control for sampling effort variation, diversity estimates were standardized using coverage-based rarefaction and extrapolation via the iNEXT framework (Chao et al., 2021). Sample coverage, defined as the proportion of total species incidences represented, served as an objective measure of completeness.

We extended iNEXT with meta-analytic methods, incorporating bootstrapped confidence intervals to aggregate effect sizes across paired forests, enabling standardized comparisons across sites and treatments.

TD was quantified as the effective number of equally abundant species, PD as the effective number of equally divergent lineages, and FD as the effective number of equally distinct functional groups (or functional “species”). Expressing all metrics as species-equivalent units allowed direct comparison across diversity dimensions.

To identify the scale of treatment effects, we decomposed γ-diversity into α-diversity (average within-patch diversity) and multiplicative β-diversity (compositional turnover among patches) using the iNEXT.beta3D package (v2.0.8; Chao et al., 2023). This decomposition applies to TD, FD, and PD under the Hill number framework (q = 0, 1, 2), where diversity is expressed as the effective number of species, functional groups, or lineages, respectively.

Multiplicative β-diversity ranges from 1 (identical patches) to the number of patches (complete dissimilarity). To facilitate interpretation, we applied the 1 − S transformation, a normalized turnover metric analogous to Jaccard dissimilarity (Chao et al., 2019; Chiu et al., 2014; Lande, 1996), scaled between 0 (no turnover) and 1 (complete turnover).

All diversity metrics were calculated from incidence data (presence/absence per trap) rather than abundance to reduce bias from short-term local fluctuations. While Hill numbers were originally developed for abundance-based data, they have been adapted for incidence data (Chao et al., 2014; Colwell et al., 2012). Incidence data better capture landscape-scale occupancy and are more robust for comparing treatments across heterogeneous habitats.

All diversity estimates were standardized to a sample coverage of 0.8, a commonly used threshold balancing site inclusion with statistical robustness. Although our design included 11 paired forest districts, only 8 met this threshold and were retained for the final analysis. This ensured reliable and consistent diversity estimates across TD, FD, and PD. TD estimates for all 11 sites and diversity estimates under alternative coverage thresholds are provided in Supplementary Material.

## Results

Our final dataset comprised 13,532 hoverflies from 120 species within the family Syrphidae, with 85% of species associated with forest habitats. Notably, six species exclusively detected in the ESBC districts are red-listed in Germany.

### Positive effects of structural heterogeneity on γ-diversity

Across all experimental regions, structurally enriched (ESBC) districts supported a higher γ-diversity of hoverflies than homogeneous (control) districts (Fig. 2). This positive effect was consistent across diversity dimensions and Hill numbers (q = 0, 1, 2), with the strongest responses observed for TD. The strongest increase in γ-diversity at q = 0 indicates that rare species, lineages, or functional groups benefited most from enhanced forest structure, while the smaller gains at q = 1 and q = 2 suggest that the benefits were less pronounced for common and dominant ones.

**Figure 2.**
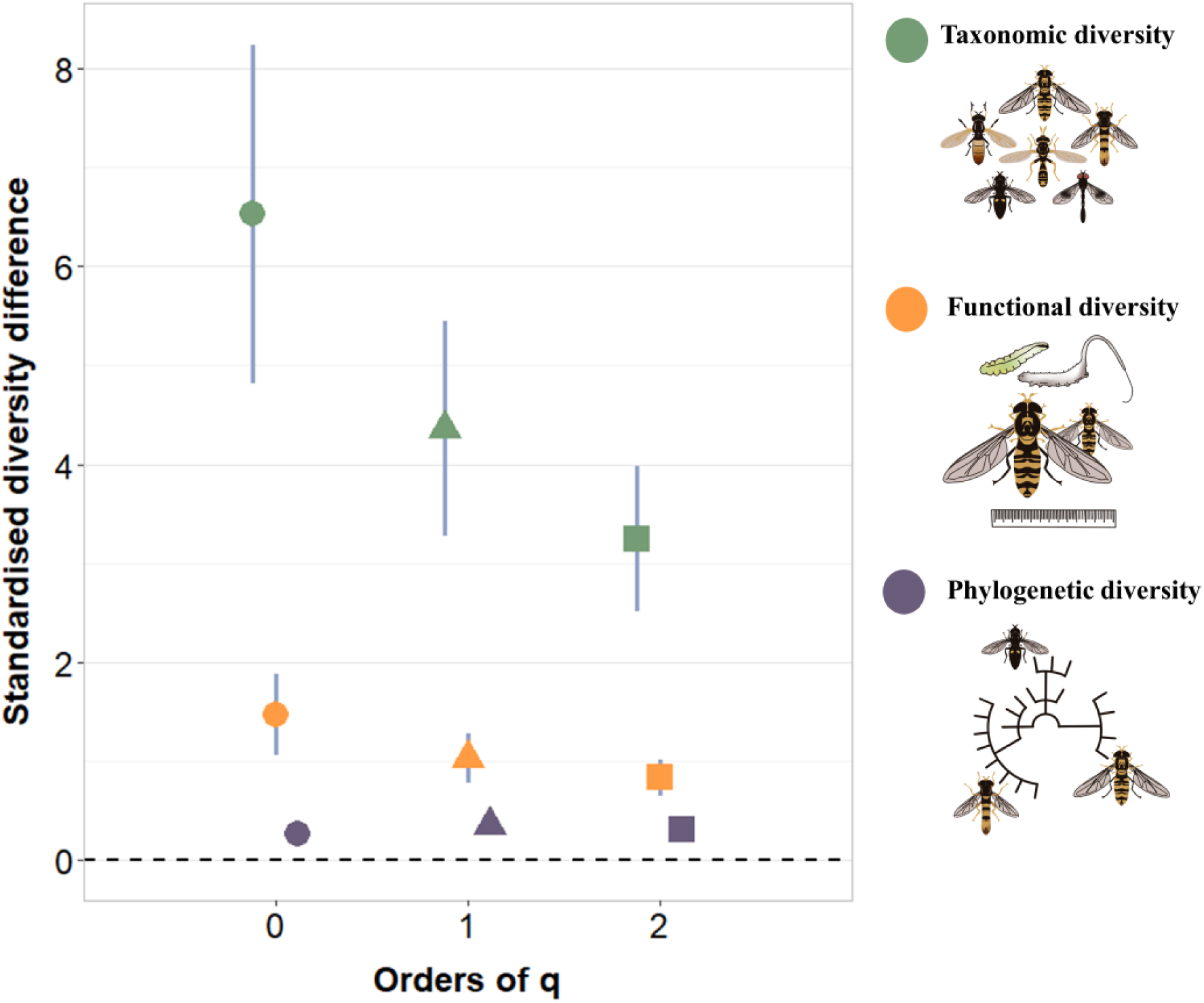
Effects of landscape heterogeneity on taxonomic, functional and phylogenetic gamma diversity across diversity orders (Order q). Points represent the mean difference between ESBC and Control forests for each diversity type, with error bars showing lower and upper confidence limits. Colours indicate diversity types (taxonomic: green, functional: orange, phylogenetic: purple). The shapes differentiate between the diversity orders (q = 0: circle, q = 1: triangle, q = 2: square). All diversity estimates were standardized to a sample coverage of 0.8.

Overall, enhanced structural complexity increased hoverfly diversity at both the γ- and α-levels across all experimental regions. This trend was consistent across different diversity dimensions and orders. In contrast, β-diversity showed a more nuanced response, with moderate increases in TD and FD, but variable or weak patterns for PD. (Fig. 3)

**Figure 3.**
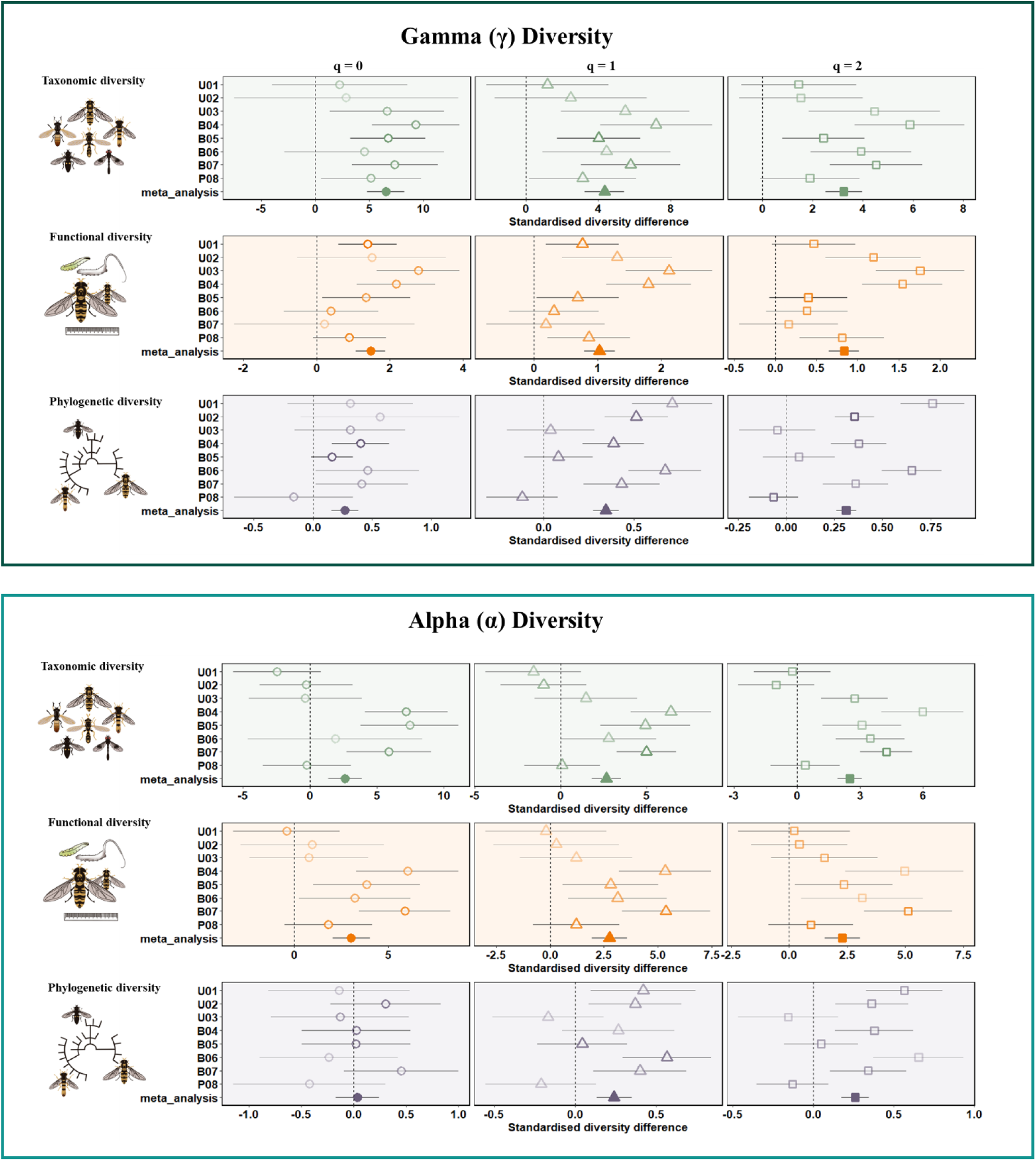
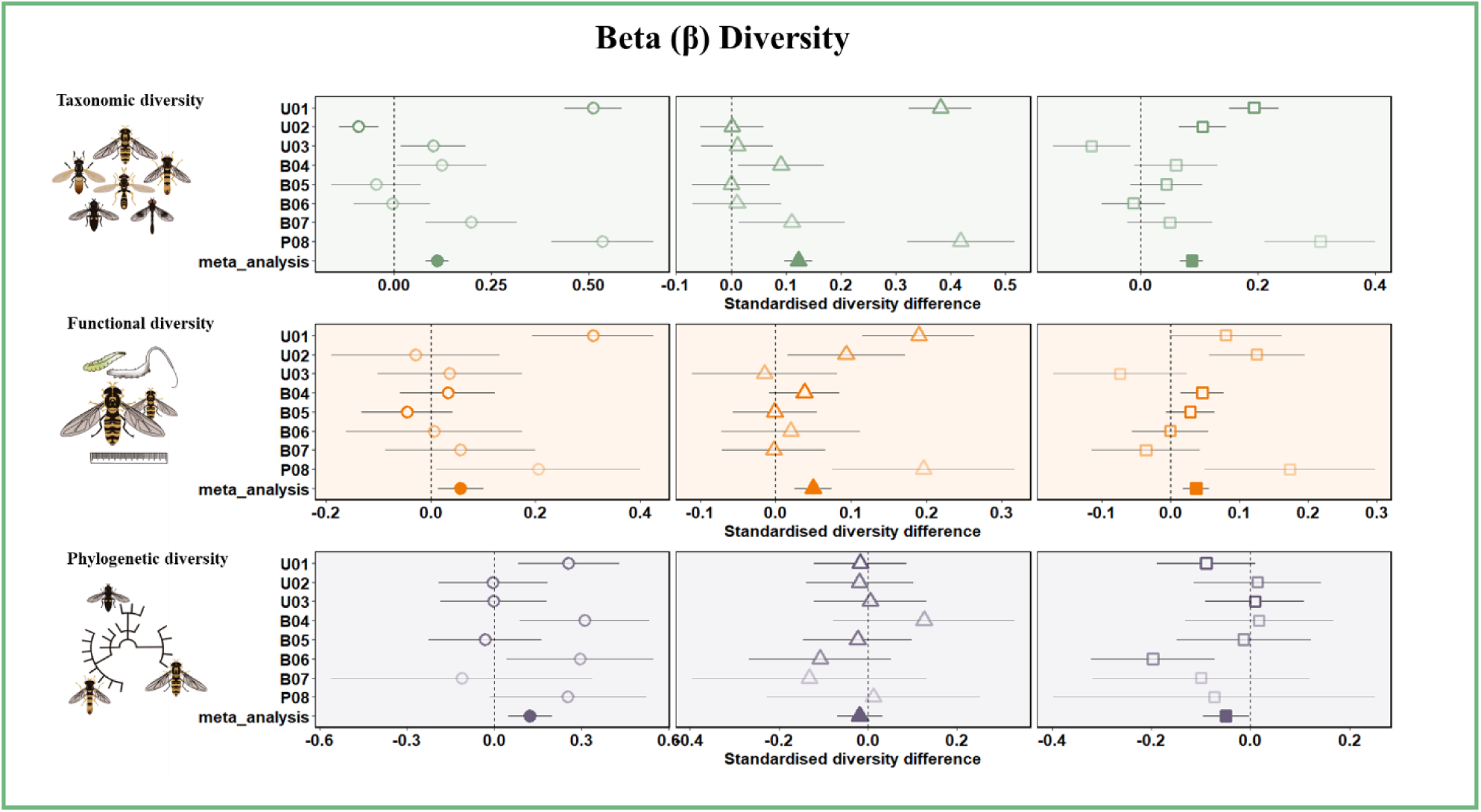
Differences in (a) taxonomic diversity, (b) functional diversity, and (c) phylogenetic diversity between ESBC and Control, shown separately for γ-diversity (gamma), α-diversity (alpha), and β-diversity (beta 1-S). Each diversity component is presented for single forest districts (open symbols) and across forest districts (meta-analysis, solid color filled symbols) for different Hill numbers (q = 0, q = 1, q = 2). Error bars indicate the 95% lower confidence limit (LCL) and the upper confidence limit (UCL). Different symbols indicate Hill numbers (circle: q = 0, triangle: q = 1, square: q = 2). Transparency levels represent group weights, with lower transparency corresponding to lower weights. Vertical dashed lines at zero denote no difference. All diversity estimates were standardized to a sample coverage of 0.8.

### α- and β-diversity: mechanisms behind γ-gains

To disentangle the mechanisms driving γ-increases, we examined α- and β-components separately (Fig. 3). In nearly all regions, gains in γ-diversity were primarily driven by higher α-diversity, suggesting that individual patches in ESBC districts supported more species, functional traits, and evolutionary lineages. This indicates that increased within-patch complexity and among-patch heterogeneity enhanced local habitat suitability for hoverflies.

In contrast, β-diversity showed more moderate and variable responses. For TD and FD, ESBC generally increased turnover among patches, consistent with a species sorting mechanism, where structurally distinct patches harbour different hoverfly assemblages. However, PD β-diversity responded weakly or even declined at higher diversity orders, suggesting that structurally heterogeneous patches hosted similar phylogenetic lineages, even when species richness increased.

Importantly, the magnitude and relative contribution of α- and β-responses differed among regions. In some districts, γ-diversity gains were almost exclusively due to local α-increases, whereas in others, moderate β-diversity gains indicated a role for landscape-level complementarity. This regional variability highlights the context dependency of how structural complexity shapes different biodiversity components across forest landscapes (Fig.3).

## Discussion

Our multiscale experimental study of hoverfly diversity reveals how spatial scale and structural forest heterogeneity influence taxonomic, functional, and phylogenetic diversity, with direct implications for forest management. We found a consistent increase in γ-diversity in heterogeneous compared to homogeneous landscapes, mainly driven by higher α-diversity within patches. This indicates that enhancing structural heterogeneity through diverse interventions effectively promotes local species richness, scaling up to greater landscape diversity. Similar patterns were observed by Rothacher (et al., 2025), where insect α-diversity was higher in canopy gaps. In contrast, evidence for species-sorting effects leading to higher β-diversity was lower and more variable, suggesting stronger regional environmental effects.

### Enhanced structural complexity scales up to higher landscape diversity

Across all diversity dimensions, γ-diversity increased primarily due to α-diversity gains under ESBC treatments, most pronounced for TD, where species richness per patch was consistently higher. These findings support the habitat heterogeneity hypothesis: increased structural complexity creates niches, boosting local richness (Cramer & Willig, 2005; Tews et al., 2004). Interventions like canopy gaps and deadwood retention benefit hoverfly species associated with open or saproxylic habitats (Achury et al., 2023; Meyer et al., 2009), consistent with studies reporting higher insect α-diversity in open canopies (Knuff et al., 2020).

The dominance of α-driven responses suggests that small-scale structural enrichment creates diverse microhabitats, enabling local niche differentiation and supporting hoverfly diversity. This matches findings for bees and butterflies in agricultural landscapes, where local habitat quality outweighs spatial configuration (Ballare et al., 2019; Bottero et al., 2023). However, the relatively low general contribution of β-diversity may also indicate that the structural complexity enhancement across patches created similar microhabitats (Bracewell et al., 2018; Cunha et al., 2019). Hoverflies appear to respond more to general features such as increased light availability and deadwood enrichment rather than specific configurations (e.g. standing vs. lying deadwood). The different treatments we implemented may have been too ecologically similar from the perspective of hoverflies, leading to functionally convergent communities. Consequently, we observe increases in α-diversity across enriched patches, but relatively low compositional dissimilarity among them.

Nonetheless, in some regions, γ-diversity was shaped by both α- and β-diversity. This highlights that while α-driven gains dominate, species turnover matters where regional differences or environmental gradients are pronounced (Ganuza et al., 2022), emphasizing the need to preserve heterogeneity both within and between patches.

### Landscape-level effects and mass effects across enhanced patches

Beyond the direct effects of enhancement, we also observed that increased γ-diversity in ESBC landscapes stemmed from cumulative gains across multiple enhanced patches. This may reflect mass effects, where species from one patch disperse into nearby patches, increasing local diversity, or by differences in habitat complementarity between enriched patches (Cadotte, 2006). Such dynamics highlight how landscape-level habitat improvements can boost α-diversity even without strong turnover between patches.

This phenomenon is particularly relevant for mobile insect groups like hoverflies, which are known for their dispersal capacity and ecological flexibility (Meyer et al., 2009; Moquet et al., 2018). Adult hoverflies exploit a wide range of floral resources, while larval stages utilise diverse microhabitats, including decaying wood and aquatic environments (Moquet et al., 2018). This flexibility likely enhances their responsiveness to structural enrichment at both local and landscape scales. Our observation of six red-listed species from Germany found exclusively in ESBC patches further underscores the conservation potential of these practices (Saure, 2018).

### Species sorting and spatial turnover

Although β-diversity generally contributed less to γ-diversity than α-diversity, in a few study regions, we observed a deviation from this pattern. In these regions, increased γ-diversity emerged despite relatively stable α-values, suggesting that spatial turnover played a larger role. This implies that in certain contexts, particularly where environmental filters differ sharply between patches or where natural disturbance regimes operate, species sorting can shape community composition (Questad & Foster, 2008).

These findings are consistent with studies emphasizing the importance of spatial heterogeneity for β-diversity, especially in systems with strong microclimatic or structural contrasts (Heino et al., 2015; Hendrickx et al., 2009; Herrault et al., 2016). While our interventions generate forest gaps and introduce deadwood, they still occur within the same general habitat type (i.e. temperate forests). Thus, compared to studies with broader habitat contrasts (e.g. inclusion of grasslands or agricultural edges), our ability to promote β-diversity may be constrained. Nevertheless, in some districts, particularly those that likely exhibited greater natural heterogeneity or more pronounced microclimatic differences, hoverfly communities responded with higher compositional dissimilarity, indicating a potential for species sorting given sufficient environmental variability.

### Divergent responses among taxonomic, functional, and phylogenetic diversity

Despite consistent TD increases, FD and PD responses were more subdued, likely due to trait redundancy and phylogenetic clustering within hoverflies (Cadotte et al., 2011). Overlapping larval habitat preferences limit functional and phylogenetic diversification (Moquet et al., 2018). Thus, increased species numbers may not translate to greater ecosystem functionality or evolutionary distinctiveness (LaRue et al., 2019). Increases were strongest at q = 0, indicating rare species benefit most, underscoring the importance of evaluating multiple biodiversity dimensions when assessing the ecological benefits of forest interventions.

### Management implications

Our results have direct implications for forest biodiversity conservation. The strong α-diversity response to ESBC suggests that local enrichment practices, such as increasing deadwood, promoting uneven-aged stands, and creating canopy gaps, are effective for supporting hoverfly diversity. These strategies align with calls to prioritize structural complexity over uniform production forests (Chisholm & Dutta Gupta, 2023; Duflot et al., 2022).

Moreover, the relatively smaller and more variable contribution of β-diversity implies that, while structural enrichment generally benefits species richness at the patch level, it may not always lead to higher community heterogeneity, that is, to differences in species composition among patches (Burrascano et al., 2018; Harrison et al., 2006). This could be due to insufficient ecological contrast between enriched patches; more pronounced interventions, such as larger canopy gaps or the promotion of early-successional habitats, may be necessary to foster species turnover and increase β-diversity. Although α-diversity effects dominated, some sites benefited from heterogeneity, suggesting that under certain conditions, structural contrast can support distinct communities. Even though the mechanisms remain unclear, managers should consider spatially differentiated enrichment strategies, including variation in gap size, canopy cover, and deadwood types, to enhance both α- and β-diversity (Larrieu et al., 2014; Uhl et al., 2022).

Incorporating scale-explicit management plans is particularly crucial. While our findings support small-scale interventions, enhancing patch-level richness, they also suggest that broader-scale heterogeneity may be overlooked without variation between treatments or across patches. Therefore, conservation planning should emphasize mosaic landscapes (Hardt et al., 2013; Vimal et al., 2017) with a mix of enrichment intensities and configurations (Vasiljević et al., 2024) to maintain both local and landscape biodiversity.

### A unique contribution to forest biodiversity research

To our knowledge, this is among the first studies to examine taxonomic, functional, and phylogenetic diversity responses simultaneously across α-, β-, and γ-levels in a landscape-scale forest experiment. By providing this multidimensional view, our study contributes valuable evidence that local-scale modifications can be a key lever for enhancing insect diversity in forests, while also emphasizing the need for spatial planning to address community turnover. By replicating interventions across multiple forest landscapes and employing a meta-analytical approach, our findings are robust and broadly applicable beyond individual study regions.

While our study provides compelling evidence for the role of structural heterogeneity in enhancing biodiversity, several limitations should be acknowledged. First, hoverflies, though ecologically significant, may not fully represent other communities and taxa with more specific habitat requirements. Second, while our experimental design captured short-term responses, the long-term effects of ESBC on biodiversity and ecosystem functioning remain uncertain. Long-term studies tracking species persistence and population dynamics will be critical for understanding the durability of these effects. Lastly, the extent to which ESBC interacts with other environmental factors, such as landscape connectivity and climate variability, warrants further investigation.

## Conclusion

Given the wide geographical range of our sampling across Germany, our results likely reflect generalizable patterns relevant to Central European production forests. We show that structurally heterogeneous forest landscapes support higher γ-diversity of hoverflies compared to homogeneous ones, underlining the ecological value of fine-scale structural interventions.

By incorporating β-heterogeneity enhancement into forest management strategies, our findings offer actionable solutions to conserve biodiversity and maintain essential ecosystem functions, such as pollination and natural pest control, in production forests. More broadly, increasing structural heterogeneity at the landscape level can contribute to integrated land-use strategies aimed at sustaining biodiversity, supporting ecosystem services, and enhancing the climate resilience of multifunctional landscapes.

## Data availability statement

If accepted, the data will be archived in Zenodo (https://zenodo.org), an open-access repository. The DOI and access details will be provided upon publication.

## Acknowledgements

We thank the technical staff for maintaining the field site and the many student helpers for their support on the experimental patches. We are grateful to Nikki Sauer and Gisela Merkel-Wallner for hoverfly species identification. BETA-FOR is funded by the Deutsche Forschungsgemeinschaft (DFG, German Research Foundation, FOR 5375) – 459717468, with additional support from the Julius-Maximilians-Universität Würzburg (JMU).

## Author contributions

Clàudia Massó Estaje, Alice Claßen and Ingolf Steffan-Dewenter conceived the ideas and designed the methodology; Clàudia Massó Estaje collected the data; Clàudia Massó Estaje, Oliver Mitesser and Julia Rothacher analysed the data; Clàudia Massó Estaje, Alice Claßen and Ingolf Steffan-Dewenter led the writing of the manuscript. All authors contributed critically to the drafts and gave final approval for publication.

## References

Achury, R., Staab, M., Blüthgen, N., & Weisser, W. W. (2023). Forest gaps increase true bug diversity by recruiting open land species. Oecologia, 202(2), 299–312. 10.1007/s00442-023-05392-z

Ballare, K. M., Neff, J. L., Ruppel, R., & Jha, S. (2019). Multi-scalar drivers of biodiversity: Local management mediates wild bee community response to regional urbanization. Ecological Applications, 29(3), e01869. 10.1002/eap.1869

Barnes, A. D., Weigelt, P., Jochum, M., Ott, D., Hodapp, D., Haneda, N. F., & Brose, U. (2016). Species richness and biomass explain spatial turnover in ecosystem functioning across tropical and temperate ecosystems. Philosophical Transactions of the Royal Society B: Biological Sciences, 371(1694), 20150279. 10.1098/rstb.2015.0279

Bottero, I., Dominik, C., Schweiger, O., Albrecht, M., Attridge, E., Brown, M. J. F., Cini, E., Costa, C., De la Rúa, P., de Miranda, J. R., Di Prisco, G., Dzul Uuh, D., Hodge, S., Ivarsson, K., Knauer, A. C., Klein, A.-M., Mänd, M., Martínez-López, V., Medrzycki, P., … Stout, J. C. (2023). Impact of landscape configuration and composition on pollinator communities across different European biogeographic regions. Frontiers in Ecology and Evolution, 11. 10.3389/fevo.2023.1128228

Bracewell, S. A., Clark, G. F., & Johnston, E. L. (2018). Habitat complexity effects on diversity and abundance differ with latitude: An experimental study over 20 degrees. Ecology, 99(9), 1964–1974. 10.1002/ecy.2408

Burrascano, S., Ripullone, F., Bernardo, L., Borghetti, M., Carli, E., Colangelo, M., Gangale, C., Gargano, D., Gentilesca, T., Luzzi, G., Passalacqua, N., Pelle, L., Rivelli, A. R., Sabatini, F. M., Schettino, A., Siclari, A., Uzunov, D., & Blasi, C. (2018). It’s a long way to the top: Plant species diversity in the transition from managed to old-growth forests. Journal of Vegetation Science, 29(1), 98–109. 10.1111/jvs.12588

Cadotte, M. W. (2006). Dispersal and Species Diversity: A Meta-Analysis. The American Naturalist, 167(6), 913–924. 10.1086/504850

Cadotte, M. W., Carscadden, K., & Mirotchnick, N. (2011). Beyond species: Functional diversity and the maintenance of ecological processes and services. Journal of Applied Ecology, 48(5), 1079–1087. 10.1111/j.1365-2664.2011.02048.x

Chao, A., Chiu, C.-H., Wu, S.-H., Huang, C.-L., & Lin, Y.-C. (2019). Comparing two classes of alpha diversities and their corresponding beta and (dis)similarity measures, with an application to the Formosan sika deer Cervus nippon taiouanus reintroduction programme. Methods in Ecology and Evolution, 10(8), 1286–1297. 10.1111/2041-210X.13233

Chao, A., Gotelli, N. J., Hsieh, T. C., Sander, E. L., Ma, K. H., Colwell, R. K., & Ellison, A. M. (2014). Rarefaction and extrapolation with Hill numbers: A framework for sampling and estimation in species diversity studies. Ecological Monographs, 84(1), 45–67. 10.1890/13-0133.1

Chao, A., Henderson, P. A., Chiu, C., Moyes, F., Hu, K., Dornelas, M., & Magurran, A. E. (2021). Measuring temporal change in alpha diversity: A framework integrating taxonomic, phylogenetic and functional diversity and the INEXT.3D standardization. Methods in Ecology and Evolution, 12(10), 1926–1940. 10.1111/2041-210X.13682

Chao, A., & Jost, L. (2012). Coverage-based rarefaction and extrapolation: Standardizing samples by completeness rather than size. Ecology, 93(12), 2533–2547. 10.1890/11-1952.1

Chao, A., Thorn, S., Chiu, C.-H., Moyes, F., Hu, K.-H., Chazdon, R. L., Wu, J., Magnago, L. F. S., Dornelas, M., Zelený, D., Colwell, R. K., & Magurran, A. E. (2023). Rarefaction and extrapolation with beta diversity under a framework of Hill numbers: The iNEXT.beta3D standardization. Ecological Monographs, 93(4), e1588. 10.1002/ecm.1588

Chisholm, R. A., & Dutta Gupta, T. (2023). A critical assessment of the biodiversity– productivity relationship in forests and implications for conservation. Oecologia, 201(4), 887–900. 10.1007/s00442-023-05363-4

Chiu, C.-H., Jost, L., & Chao, A. (2014). Phylogenetic beta diversity, similarity, and differentiation measures based on Hill numbers. Ecological Monographs, 84(1), 21–44. 10.1890/12-0960.1

Colwell, R. K., Chao, A., Gotelli, N. J., Lin, S.-Y., Mao, C. X., Chazdon, R. L., & Longino, J. T. (2012). Models and estimators linking individual-based and sample-based rarefaction, extrapolation and comparison of assemblages. Journal of Plant Ecology, 5(1), 3–21. 10.1093/jpe/rtr044

Cramer, M. J., & Willig, M. R. (2005). Habitat heterogeneity, species diversity and null models. Oikos, 108(2), 209–218. 10.1111/j.0030-1299.2005.12944.x

Cunha, E. R., Winemiller, K. O., da Silva, J. C. B., Lopes, T. M., Gomes, L. C., Thomaz, S. M., & Agostinho, A. A. (2019). α and β diversity of fishes in relation to a gradient of habitat structural complexity supports the role of environmental filtering in community assembly. Aquatic Sciences, 81(2), 38. 10.1007/s00027-019-0634-3

Dubois, S., & Chamberlain, S. (2025). bold: Interface to Bold Systems API.

Duflot, R., Fahrig, L., & Mönkkönen, M. (2022). Management diversity begets biodiversity in production forest landscapes. Biological Conservation, 268, 109514. 10.1016/j.biocon.2022.109514

Eckelt, A., Müller, J., Bense, U., Brustel, H., Bußler, H., Chittaro, Y., Cizek, L., Frei, A., Holzer, E., Kadej, M., Kahlen, M., Köhler, F., Möller, G., Mühle, H., Sanchez, A., Schaffrath, U., Schmidl, J., Smolis, A., Szallies, A., … Seibold, S. (2018). “Primeval forest relict beetles” of Central Europe: A set of 168 umbrella species for the protection of primeval forest remnants. Journal of Insect Conservation, 22(1), 15–28. 10.1007/s10841-017-0028-6

Faith, D. P. (2016). Using Phylogenetic Dissimilarities Among Sites for Biodiversity Assessments and Conservation. In R. Pellens & P. Grandcolas (Eds.), Biodiversity Conservation and Phylogenetic Systematics: Preserving our evolutionary heritage in an extinction crisis (pp. 119–139). Springer International Publishing. 10.1007/978-3-319-22461-9_7

Ganuza, C., Redlich, S., Uhler, J., Tobisch, C., Rojas-Botero, S., Peters, M. K., Zhang, J., Benjamin, C. S., Englmeier, J., Ewald, J., Fricke, U., Haensel, M., Kollmann, J., Riebl, R., Uphus, L., Müller, J., & Steffan-Dewenter, I. (2022). Interactive effects of climate and land use on pollinator diversity differ among taxa and scales. Science Advances, 8(18), eabm9359. 10.1126/sciadv.abm9359

Gonçalves-Souza, T., Chase, J. M., Haddad, N. M., Vancine, M. H., Didham, R. K., Melo, F. L. P., Aizen, M. A., Bernard, E., Chiarello, A. G., Faria, D., Gibb, H., de Lima, M. G., Magnago, L. F. S., Mariano-Neto, E., Nogueira, A. A., Nemésio, A., Passamani, M., Pinho, B. X., Rocha-Santos, L., … Sanders, N. J. (2025). Species turnover does not rescue biodiversity in fragmented landscapes. Nature, 640(8059), 702–706. 10.1038/s41586-025-08688-7

Hardt, E., dos Santos, R. F., de Pablo, C. L., de Agar, P. M., & Pereira-Silva, E. F. L. (2013). Utility of landscape mosaics and boundaries in forest conservation decision making in the Atlantic Forest of Brazil. Landscape Ecology, 28(3), 385–399. 10.1007/s10980-013-9845-5

Harrison, S., Davies, K. F., Safford, H. D., & Viers, J. H. (2006). Beta diversity and the scale-dependence of the productivity-diversity relationship: A test in the Californian serpentine flora. Journal of Ecology, 94(1), 110–117. 10.1111/j.1365-2745.2005.01078.x

Heidrich, L., Bae, S., Levick, S., Seibold, S., Weisser, W., Krzystek, P., Magdon, P., Nauss, T., Schall, P., Serebryanyk, A., Wöllauer, S., Ammer, C., Bässler, C., Doerfler, I., Fischer, M., Gossner, M. M., Heurich, M., Hothorn, T., Jung, K., … Müller, J. (2020). Heterogeneity–diversity relationships differ between and within trophic levels in temperate forests. Nature Ecology & Evolution, 4(9), 1204–1212. 10.1038/s41559-020-1245-z

Heino, J., Melo, A. S., & Bini, L. M. (2015). Reconceptualising the beta diversity-environmental heterogeneity relationship in running water systems. Freshwater Biology, 60(2), 223–235. 10.1111/fwb.12502

Hendrickx, F., Maelfait, J.-P., Desender, K., Aviron, S., Bailey, D., Diekotter, T., Lens, L., Liira, J., Schweiger, O., Speelmans, M., Vandomme, V., & Bugter, R. (2009). Pervasive effects of dispersal limitation on within- and among-community species richness in agricultural landscapes. Global Ecology and Biogeography, 18(5), 607– 616. 10.1111/j.1466-8238.2009.00473.x

Herrault, P.-A., Larrieu, L., Cordier, S., Gimmi, U., Lachat, T., Ouin, A., Sarthou, J.-P., & Sheeren, D. (2016). Combined effects of area, connectivity, history and structural heterogeneity of woodlands on the species richness of hoverflies (Diptera: Syrphidae). Landscape Ecology, 31(4), 877–893. 10.1007/s10980-015-0304-3

Hill, M. O. (1973). Diversity and evenness: A unifying notation and its consequences. Ecology, 54(2), 427–432.

Hilmers, T., Friess, N., Bässler, C., Heurich, M., Brandl, R., Pretzsch, H., Seidl, R., & Müller, J. (2018). Biodiversity along temperate forest succession. Journal of Applied Ecology, 55(6), 2756–2766. 10.1111/1365-2664.13238

Knuff, A. K., Staab, M., Frey, J., Dormann, C. F., Asbeck, T., & Klein, A.-M. (2020). Insect abundance in managed forests benefits from multi-layered vegetation. Basic and Applied Ecology, 48, 124–135. 10.1016/j.baae.2020.09.002

Kortmann, M., Chao, A., Schaefer, H. M., Blüthgen, N., Gelis, R., Tremlett, C. J., Busse, A., Püls, M., Seibold, S., Kriegel, P., Rabl, D., De La Hoz, M., Şekercioğlu, Ç. H., Schleuning, M., Feldhaar, H., Newell, F. L., Kümmet, S., Mitesser, O., Peters, M. K., & Müller, J. (2025). Sample coverage affects diversity measures of bird communities along a natural recovery gradient of abandoned agriculture in tropical lowland forests. Journal of Applied Ecology, 62(3), 480–491. 10.1111/1365-2664.14879

Lande, R. (1996). Statistics and Partitioning of Species Diversity, and Similarity among Multiple Communities. Oikos, 76(1), 5–13. 10.2307/3545743

Larrieu, L., Cabanettes, A., Gonin, P., Lachat, T., Paillet, Y., Winter, S., Bouget, C., & Deconchat, M. (2014). Deadwood and tree microhabitat dynamics in unharvested temperate mountain mixed forests: A life-cycle approach to biodiversity monitoring. Forest Ecology and Management, 334, 163–173. 10.1016/j.foreco.2014.09.007

LaRue, E. A., Hardiman, B. S., Elliott, J. M., & Fei, S. (2019). Structural diversity as a predictor of ecosystem function. Environmental Research Letters, 14(11), 114011. 10.1088/1748-9326/ab49bb

Martin, E. A., Dainese, M., Clough, Y., Báldi, A., Bommarco, R., Gagic, V., Garratt, M. P. D., Holzschuh, A., Kleijn, D., Kovács-Hostyánszki, A., Marini, L., Potts, S. G., Smith, H. G., Hassan, D. A., Albrecht, M., Andersson, G. K. S., Asís, J. D., Aviron, S., Balzan, M. V., … Steffan-Dewenter, I. (2019). The interplay of landscape composition and configuration: New pathways to manage functional biodiversity and agroecosystem services across Europe. Ecology Letters, 22(7), 1083–1094. 10.1111/ele.13265

McGill, B. J., Dornelas, M., Gotelli, N. J., & Magurran, A. E. (2015). Fifteen forms of biodiversity trend in the Anthropocene. Trends in Ecology & Evolution, 30(2), 104–113. 10.1016/j.tree.2014.11.006

Meyer, B., Jauker, F., & Steffan-Dewenter, I. (2009). Contrasting resource-dependent responses of hoverfly richness and density to landscape structure. Basic and Applied Ecology, 10(2), 178–186. 10.1016/j.baae.2008.01.001

Moquet, L., Laurent, E., Bacchetta, R., & Jacquemart, A. (2018). Conservation of hoverflies (Diptera, Syrphidae) requires complementary resources at the landscape and local scales. Insect Conservation and Diversity, 11(1), 72–87. 10.1111/icad.12245

Mori, A. S., Isbell, F., & Seidl, R. (2018). β-Diversity, Community Assembly, and Ecosystem Functioning. Trends in Ecology & Evolution, 33(7), 549–564. 10.1016/j.tree.2018.04.012

Müller, J., Mitesser, O., Cadotte, M. W., van der Plas, F., Mori, A. S., Ammer, C., Chao, A., Scherer-Lorenzen, M., Baldrian, P., Bässler, C., Biedermann, P., Cesarz, S., Claßen, A., Delory, B. M., Feldhaar, H., Fichtner, A., Hothorn, T., Kuenzer, C., Peters, M. K., … Eisenhauer, N. (2023). Enhancing the structural diversity between forest patches— A concept and real-world experiment to study biodiversity, multifunctionality and forest resilience across spatial scales. Global Change Biology, 29(6), 1437–1450. 10.1111/gcb.16564

Ódor, P., Heilmann-Clausen, J., Christensen, M., Aude, E., van Dort, K. W., Piltaver, A., Siller, I., Veerkamp, M. T., Walleyn, R., Standovár, T., van Hees, A. F. M., Kosec, J., Matočec, N., Kraigher, H., & Grebenc, T. (2006). Diversity of dead wood inhabiting fungi and bryophytes in semi-natural beech forests in Europe. Biological Conservation, 131(1), 58–71. 10.1016/j.biocon.2006.02.004

Oehri, J., Schmid, B., Schaepman-Strub, G., & Niklaus, P. A. (2017). Biodiversity promotes primary productivity and growing season lengthening at the landscape scale. Proceedings of the National Academy of Sciences, 114(38), 10160–10165. 10.1073/pnas.1703928114

Perović, D., Gámez-Virués, S., Börschig, C., Klein, A.-M., Krauss, J., Steckel, J., Rothenwöhrer, C., Erasmi, S., Tscharntke, T., & Westphal, C. (2015). Configurational landscape heterogeneity shapes functional community composition of grassland butterflies. Journal of Applied Ecology, 52(2), 505–513. 10.1111/1365-2664.12394

Popov, S., Miličić, M., Diti, I., Marko, O., Sommaggio, D., Markov, Z., & Vujić, A. (2017). Phytophagous hoverflies (Diptera: Syrphidae) as indicators of changing landscapes. Community Ecology, 18(3), 287–294. 10.1556/168.2017.18.3.7

Questad, E. J., & Foster, B. L. (2008). Coexistence through spatio-temporal heterogeneity and species sorting in grassland plant communities. Ecology Letters, 11(7), 717–726. 10.1111/j.1461-0248.2008.01186.x

R Core Team. (2023). R: A Language and Environment for Statistical Computing. R Foundation for Statistical Computing. https://www.R-project.org/

Redlich, S., Martin, E. A., Wende, B., & Steffan-Dewenter, I. (2018). Landscape heterogeneity rather than crop diversity mediates bird diversity in agricultural landscapes. PLOS ONE, 13(8), e0200438. 10.1371/journal.pone.0200438

Rothacher, J., Seidl, R., Thom, D., Kortmann, M., Chao, A., Chiu, C.-H., Heibl, C., Hothorn, T., Mitesser, O., Mori, A. S., Morinière, J., Pierick, K., Wild, C., Wild, N., & Müller, J. (2025). The impact of tree mortality and post-disturbance management on insect diversity in temperate forests: Insights from a replicated experiment. Journal of Applied Ecology, n/a(n/a). 10.1111/1365-2664.70086

Saure, C. (2018). Rote Liste und Gesamtartenliste der Schwebfliegen (Diptera: Syrphidae) von Berlin. Universitätsverlag der TU Berlin. 10.14279/DEPOSITONCE-6691

Schuldt, A., Ebeling, A., Kunz, M., Staab, M., Guimarães-Steinicke, C., Bachmann, D., Buchmann, N., Durka, W., Fichtner, A., Fornoff, F., Härdtle, W., Hertzog, L. R., Klein, A.-M., Roscher, C., Schaller, J., Von Oheimb, G., Weigelt, A., Weisser, W., Wirth, C., … Eisenhauer, N. (2019). Multiple plant diversity components drive consumer communities across ecosystems. Nature Communications, 10(1), 1460. 10.1038/s41467-019-09448-8

Sobek, S., GOßNER, M. M., Scherber, C., Steffan-Dewenter, I., & Tscharntke, T. (2009). Tree diversity drives abundance and spatiotemporal β-diversity of true bugs (Heteroptera). Ecological Entomology, 34(6), 772–782. 10.1111/j.1365-2311.2009.01132.x

Speight, M., Speight, M., Castella, E., Sarthou, J.-P., & Cédric, V. (2021). StN KEY FOR THE IDENTIFICATION OF THE GENERA OF EUROPEAN SYRPHIDAE 2020.

Tews, J., Brose, U., Grimm, V., Tielbörger, K., Wichmann, M. C., Schwager, M., & Jeltsch, F. (2004). Animal species diversity driven by habitat heterogeneity/diversity: The importance of keystone structures. Journal of Biogeography, 31(1), 79–92. 10.1046/j.0305-0270.2003.00994.x

Thomsen, M. S., Altieri, A. H., Angelini, C., Bishop, M. J., Bulleri, F., Farhan, R., Frühling, V. M. M., Gribben, P. E., Harrison, S. B., He, Q., Klinghardt, M., Langeneck, J., Lanham, B. S., Mondardini, L., Mulders, Y., Oleksyn, S., Ramus, A. P., Schiel, D. R., Schneider, T., … Zotz, G. (2022). Heterogeneity within and among co-occurring foundation species increases biodiversity. Nature Communications, 13(1), 581. 10.1038/s41467-022-28194-y

Tscharntke, T., Tylianakis, J. M., Rand, T. A., Didham, R. K., Fahrig, L., Batáry, P., Bengtsson, J., Clough, Y., Crist, T. O., Dormann, C. F., Ewers, R. M., Fründ, J., Holt, R. D., Holzschuh, A., Klein, A. M., Kleijn, D., Kremen, C., Landis, D. A., Laurance, W., … Westphal, C. (2012). Landscape moderation of biodiversity patterns and processes—Eight hypotheses. Biological Reviews, 87(3), 661–685. 10.1111/j.1469-185X.2011.00216.x

Uhl, B., Krah, F.-S., Baldrian, P., Brandl, R., Hagge, J., Müller, J., Thorn, S., Vojtech, T., & Bässler, C. (2022). Snags, logs, stumps, and microclimate as tools optimizing deadwood enrichment for forest biodiversity. Biological Conservation, 270, 109569. 10.1016/j.biocon.2022.109569

Uhl, B., Schall, P., & Bässler, C. (2024). Achieving structural heterogeneity and high multi-taxon biodiversity in managed forest ecosystems: A European review. Biodiversity and Conservation. 10.1007/s10531-024-02878-x

Vasiljević, N., Mitrović, S., Devetaković, J., & Pešić, M. (2024). Landscape approach to Forest landscape restoration (FLR): Case study of Surčin minicipality. REFORESTA, 18, Article 18. 10.21750/REFOR.18.05.121

Vimal, R., Fonderflick, J., Thompson, J. D., Pluvinet, P., Debussche, M., Cheylan, M., Géniez, P., Mathevet, R., Acquarone, A., & Lepart, J. (2017). Integrating habitat diversity into species conservation in the Mediterranean mosaic landscape. Basic and Applied Ecology, 22, 36–43. 10.1016/j.baae.2017.07.001

Wright, E. S. (2016). Using DECIPHER v2.0 to Analyze Big Biological Sequence Data in R. The R Journal, 8(1), 352–359.

