## Supplementary Material for "Experimental enhancement of structural heterogeneity in forest landscapes promotes multidimensional hoverfly diversity"

**Supplementary Method**

For four species (Eupeodes lapponicus, Heringia latitarsis, Heringia pubescens, and Heringia vitripennis) sequences were not yet incorporated in the BOLD barcoding library. We thus retrieved sequences identified to genus level for these taxa. To ensure geographical relevance, we restricted these searches to congeners sampled in Germany. In cases where multiple sequences were available for the same species, only the first sequence was retrieved.

Sequence data were saved in a FASTA file, and gaps were removed prior to analysis.

### Supplementary Figures

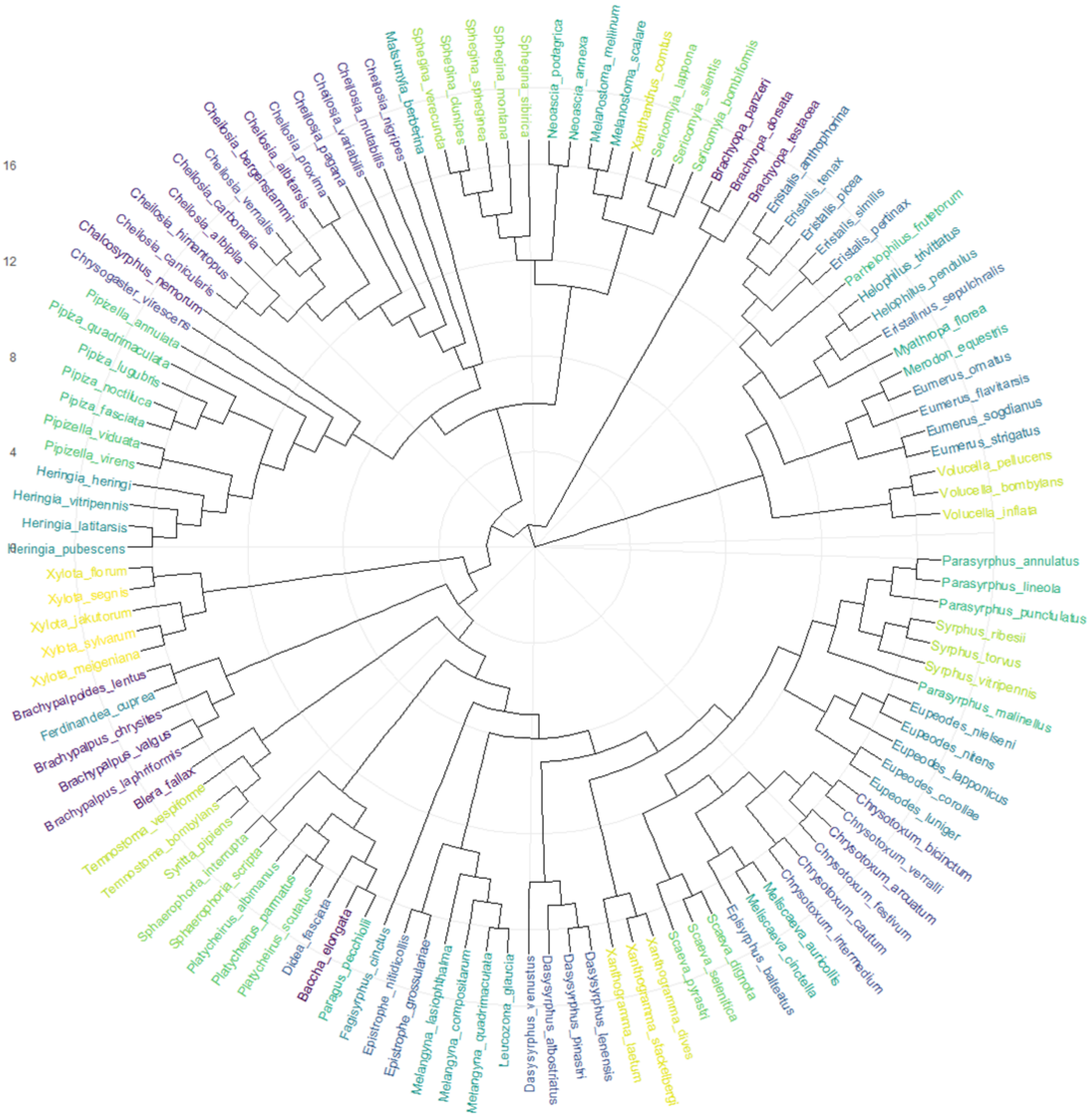


*Figure S1. Phylogenetic tree of the hoverfly species sampled in the BETA-FOR experiment. The tree illustrates the evolutionary relationships among species, constructed using the maximum likelihood method based on COI-5P sequence data from the Barcode of Life Data System (BOLD). Species names are shown at the tips, and branch lengths represent genetic divergence. This tree was used to calculate phylogenetic diversity (PD) metrics in the study.*

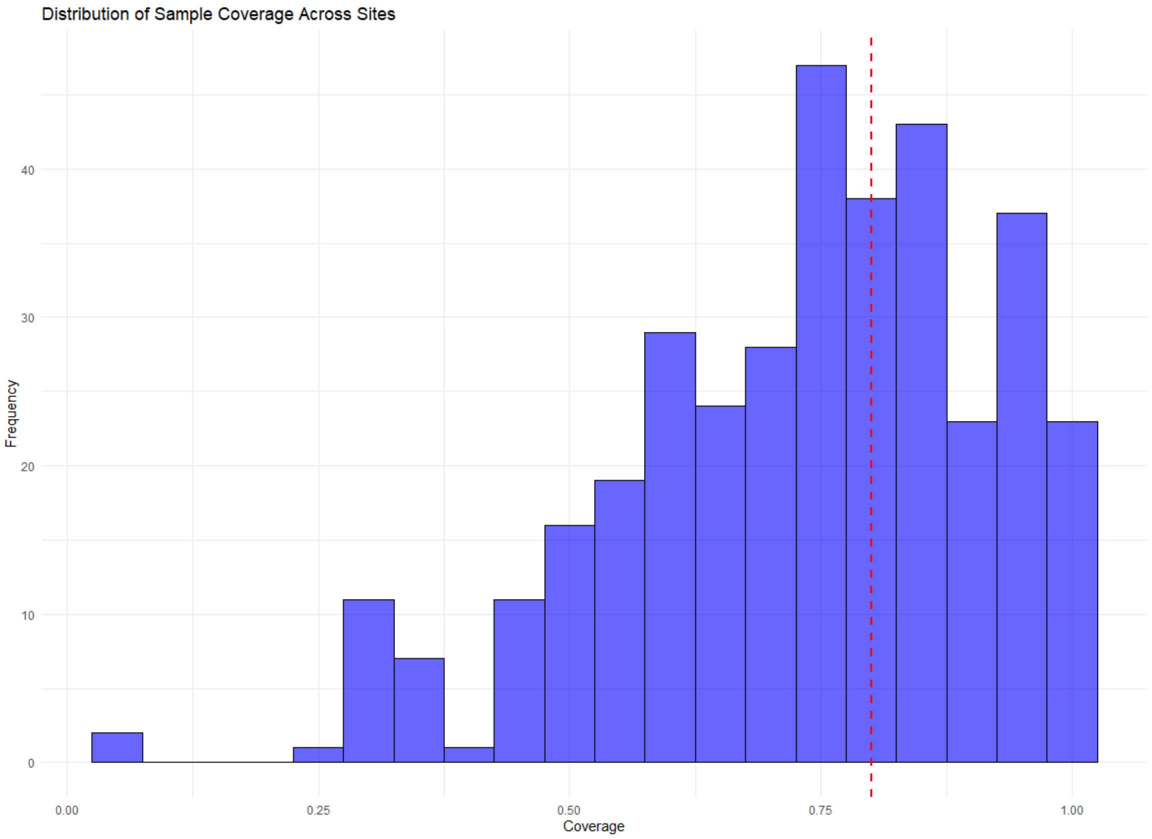


*Figure S2. Distribution of sample coverage across sites in the BETA-FOR experiment. The histogram shows the range of estimated coverage values (Cmin and Cmax) for incidence-based diversity calculations. The red line indicates the standardized coverage threshold of 0.8 used for rarefaction and extrapolation analyses.*

Taxonomic diversity – 0.8

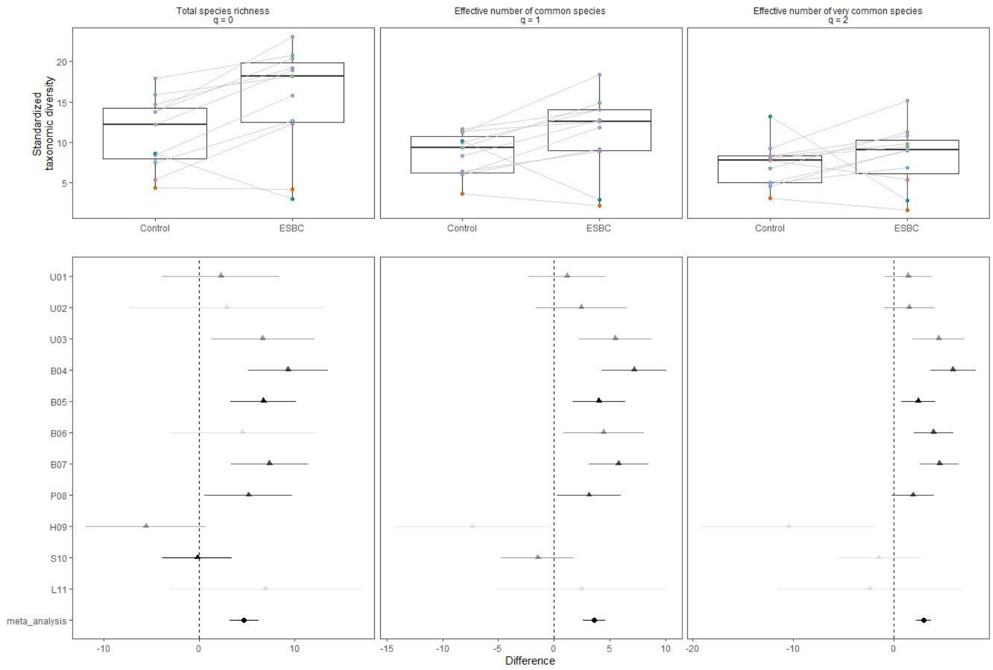


*Figure S3. Differences in taxonomic gamma diversity between ESBC and Control at a sample coverage of 0.8. Each diversity component is presented for single forest districts (triangle symbols) and across forest districts (meta-analysis, circle symbols) for different Hill numbers (q = 0, q = 1, q = 2). Error bars indicate the 95% lower confidence limit (LCL) and the upper confidence limit (UCL). Transparency levels represent group weights, with lower transparency corresponding to lower weights. Vertical dashed lines at zero denote no difference.*

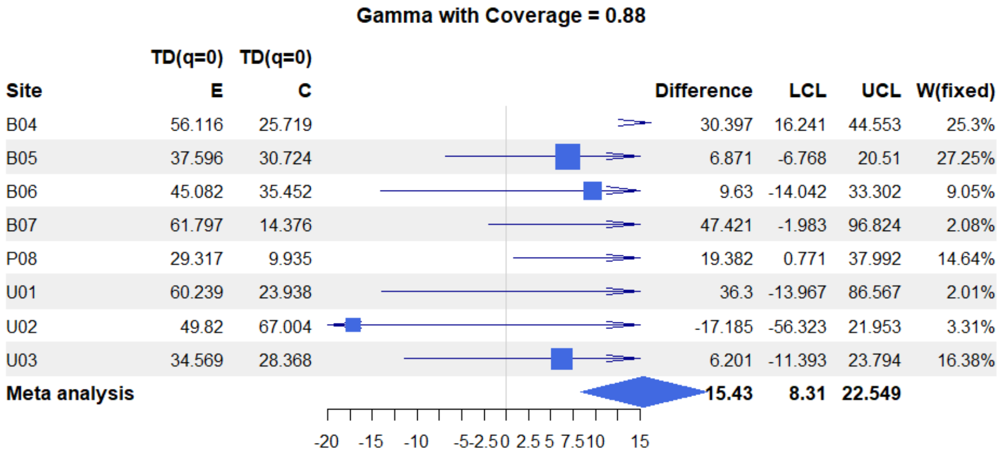


| Site | TD(q=0) E | TD(q=0) C | Difference | LCL | UCL | W(fixed) |
|---|---|---|---|---|---|---|
| B04 | 56.116 | 25.719 | 30.397 | 16.241 | 44.553 | 25.3% |
| B05 | 37.596 | 30.724 | 6.871 | -6.768 | 20.51 | 27.25% |
| B06 | 45.082 | 35.452 | 9.63 | -14.042 | 33.302 | 9.05% |
| B07 | 61.797 | 14.376 | 47.421 | -1.983 | 96.824 | 2.08% |
| P08 | 29.317 | 9.935 | 19.382 | 0.771 | 37.992 | 14.64% |
| U01 | 60.239 | 23.938 | 36.3 | -13.967 | 86.567 | 2.01% |
| U02 | 49.82 | 67.004 | -17.185 | -56.323 | 21.953 | 3.31% |
| U03 | 34.569 | 28.368 | 6.201 | -11.393 | 23.794 | 16.38% |
| **Meta analysis** | | | **15.43** | **8.31** | **22.549** | |

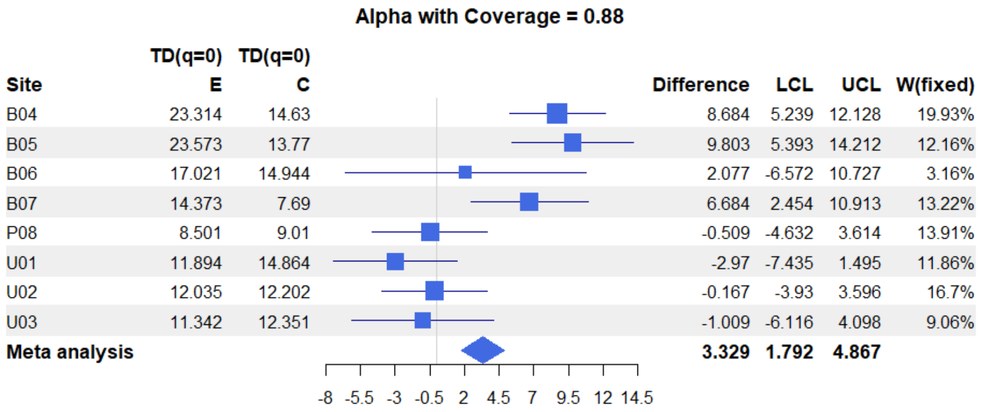


| Site | TD(q=0) E | TD(q=0) C | Difference | LCL | UCL | W(fixed) |
|---|---|---|---|---|---|---|
| B04 | 23.314 | 14.63 | 8.684 | 5.239 | 12.128 | 19.93% |
| B05 | 23.573 | 13.77 | 9.803 | 5.393 | 14.212 | 12.16% |
| B06 | 17.021 | 14.944 | 2.077 | -6.572 | 10.727 | 3.16% |
| B07 | 14.373 | 7.69 | 6.684 | 2.454 | 10.913 | 13.22% |
| P08 | 8.501 | 9.01 | -0.509 | -4.632 | 3.614 | 13.91% |
| U01 | 11.894 | 14.864 | -2.97 | -7.435 | 1.495 | 11.86% |
| U02 | 12.035 | 12.202 | -0.167 | -3.93 | 3.596 | 16.7% |
| U03 | 11.342 | 12.351 | -1.009 | -6.116 | 4.098 | 9.06% |
| **Meta analysis** | | | **3.329** | **1.792** | **4.867** | |

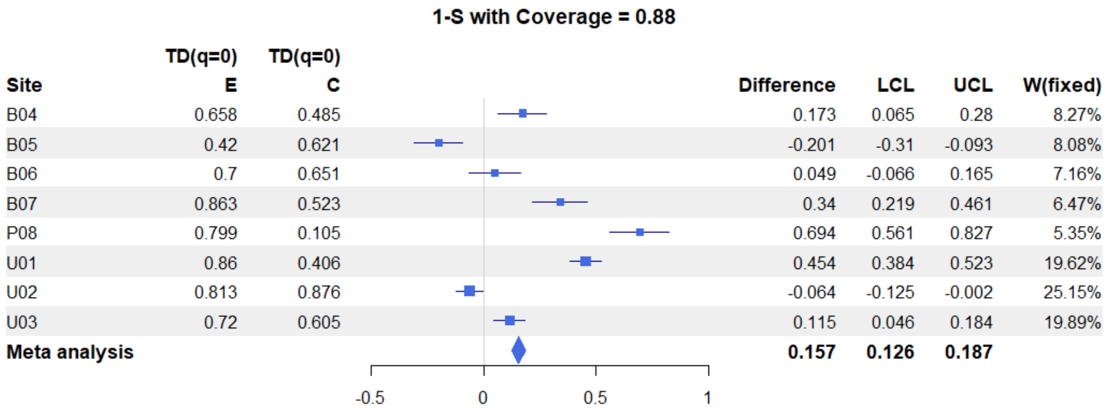


| Site | TD(q=0) E | TD(q=0) C | Difference | LCL | UCL | W(fixed) |
|---|---|---|---|---|---|---|
| B04 | 0.658 | 0.485 | 0.173 | 0.065 | 0.28 | 8.27% |
| B05 | 0.42 | 0.621 | -0.201 | -0.31 | -0.093 | 8.08% |
| B06 | 0.7 | 0.651 | 0.049 | -0.066 | 0.165 | 7.16% |
| B07 | 0.863 | 0.523 | 0.34 | 0.219 | 0.461 | 6.47% |
| P08 | 0.799 | 0.105 | 0.694 | 0.561 | 0.827 | 5.35% |
| U01 | 0.86 | 0.406 | 0.454 | 0.384 | 0.523 | 19.62% |
| U02 | 0.813 | 0.876 | -0.064 | -0.125 | -0.002 | 25.15% |
| U03 | 0.72 | 0.605 | 0.115 | 0.046 | 0.184 | 19.89% |
| **Meta analysis** | | | **0.157** | **0.126** | **0.187** | |

*Figure S4. Differences in taxonomic gamma, alpha and beta diversity between ESBC and Control at a sample coverage of 0.88. Each diversity component is presented for single forest districts (square symbols) and across forest districts (meta-analysis, diamond symbols) for q = 0. Error bars indicate the 95% lower confidence limit (LCL) and the upper confidence limit (UCL). Symbol sizes represent group weights, with larger symbols indicating higher weights in the meta-analysis. Vertical lines at zero denote no difference.*

**Gamma with Coverage = 0.721**

| Site | TD(q=0) E | TD(q=0) C | Difference | LCL | UCL | W(fixed) |
|---|---|---|---|---|---|---|
| B04 | 34.128 | 18.923 | 15.204 | 9.477 | 20.932 | 20.58% |
| B05 | 25.296 | 16.772 | 8.525 | 4.086 | 12.963 | 34.28% |
| B06 | 27.191 | 21.786 | 5.406 | -10.097 | 20.908 | 2.81% |
| B07 | 22.816 | 11.08 | 11.736 | 2.347 | 21.125 | 7.66% |
| P08 | 14.763 | 8.13 | 6.632 | 0.803 | 12.462 | 19.87% |
| U01 | 40.127 | 16.755 | 23.372 | 4.515 | 42.229 | 1.9% |
| U02 | 29.966 | 36.059 | -6.094 | -41.776 | 29.589 | 0.53% |
| U03 | 23.34 | 19.635 | 3.704 | -3.682 | 11.091 | 12.37% |
| **Meta analysis** | | | **9.29** | **6.691** | **11.888** | |

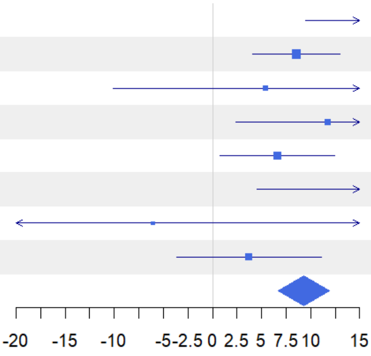


**Alpha with Coverage = 0.721**

| Site | TD(q=0) E | TD(q=0) C | Difference | LCL | UCL | W(fixed) |
|---|---|---|---|---|---|---|
| B04 | 16.892 | 11.178 | 5.714 | 3.157 | 8.271 | 15.72% |
| B05 | 15.362 | 10.431 | 4.931 | 1.624 | 8.239 | 9.39% |
| B06 | 11.928 | 10.266 | 1.661 | -3.146 | 6.469 | 4.45% |
| B07 | 10.883 | 5.779 | 5.104 | 3.119 | 7.09 | 26.07% |
| P08 | 6.764 | 6.74 | 0.024 | -2.653 | 2.701 | 14.34% |
| U01 | 9.243 | 11.229 | -1.985 | -5.223 | 1.253 | 9.8% |
| U02 | 9.429 | 9.838 | -0.409 | -3.35 | 2.532 | 11.88% |
| U03 | 9.389 | 9.093 | 0.297 | -3.213 | 3.806 | 8.34% |
| **Meta analysis** | | | **2.551** | **1.538** | **3.565** | |

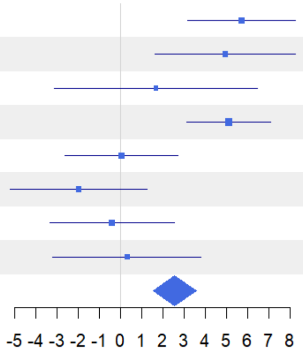


**Beta with Coverage = 0.721**

| Site | TD(q=0) E | TD(q=0) C | Difference | LCL | UCL | W(fixed) |
|---|---|---|---|---|---|---|
| B04 | 2.02 | 1.693 | 0.327 | -0.555 | 1.21 | 40.67% |
| B05 | 1.647 | 1.608 | 0.039 | -0.993 | 1.071 | 29.73% |
| B06 | 2.28 | 2.122 | 0.158 | -2.708 | 3.023 | 3.86% |
| B07 | 2.097 | 1.917 | 0.179 | -1.81 | 2.168 | 8.01% |
| P08 | 2.182 | 1.206 | 0.976 | -1.504 | 3.457 | 5.15% |
| U01 | 4.341 | 1.492 | 2.849 | -0.048 | 5.746 | 3.77% |
| U02 | 3.178 | 3.665 | -0.487 | -2.808 | 1.834 | 5.88% |
| U03 | 2.486 | 2.159 | 0.326 | -2.966 | 3.619 | 2.92% |
| **Meta analysis** | | | **0.304** | **-0.259** | **0.867** | |

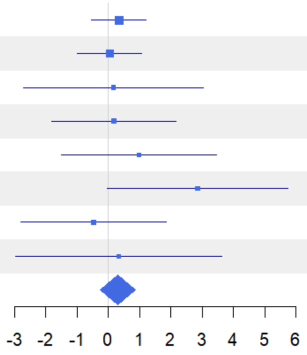


*Figure S5. Differences in taxonomic gamma, alpha and beta diversity between ESBC and Control at a sample coverage of 0.721. Each diversity component is presented for single forest districts (square symbols) and across forest districts (meta-analysis, diamond symbols) for q = 0. Error bars indicate the 95% lower confidence limit (LCL) and the upper confidence limit (UCL). Symbol sizes represent group weights, with larger symbols indicating higher weights in the meta-analysis. Vertical lines at zero denote no difference.*

### Supplementary Tables

*Table S1. Results from the gamma meta-analysis comparing taxonomic, functional, and phylogenetic diversity between ESBC and Control forests across study sites. The table includes effect size estimates (Difference), standard errors (SE), and confidence intervals (LCL, UCL) for each diversity order (Order q). The weight fixed column represents the relative contribution of each site to the overall meta-analysis estimate.*

| Site | Difference | SE | LCL | UCL | Order q | Diversity | weight fixed |
|---|---|---|---|---|---|---|---|
| U03 | 6.661722 | 2.698087 | 1.373568 | 11.949876 | 0 | TD | 10.45856 |
| U01 | 2.270497 | 3.209456 | -4.019922 | 8.560917 | 0 | TD | 7.39130 |
| U02 | 2.871914 | 5.298726 | -7.513399 | 13.257228 | 0 | TD | 2.71169 |
| B04 | 9.318544 | 2.073410 | 5.254734 | 13.382353 | 0 | TD | 17.70980 |
| B05 | 6.729671 | 1.773790 | 3.253106 | 10.206237 | 0 | TD | 24.19800 |
| B06 | 4.547875 | 3.778581 | -2.858007 | 11.953759 | 0 | TD | 5.33244 |
| B07 | 7.362218 | 2.028173 | 3.387070 | 11.337365 | 0 | TD | 18.50861 |
| P08 | 5.159170 | 2.358290 | 0.537005 | 9.781336 | 0 | TD | 13.68955 |
| meta-analysis | 6.532585 | 0.872553 | 4.822411 | 8.242759 | 0 | TD | 100 |
| U03 | 5.496108 | 1.811364 | 1.945899 | 9.046317 | 1 | TD | 9.41000 |
| U01 | 1.186908 | 1.717472 | -2.179276 | 4.553093 | 1 | TD | 10.46698 |
| U02 | 2.470734 | 2.150110 | -1.743405 | 6.684874 | 1 | TD | 6.67851 |
| B04 | 7.203530 | 1.581095 | 4.104640 | 10.302420 | 1 | TD | 12.35052 |
| B05 | 4.036809 | 1.176782 | 1.730359 | 6.343260 | 1 | TD | 22.29509 |
| B06 | 4.459270 | 1.804426 | 0.922660 | 7.995880 | 1 | TD | 9.48250 |
| B07 | 5.790025 | 1.401531 | 3.043074 | 8.536975 | 1 | TD | 15.71794 |
| P08 | 3.141656 | 1.506802 | 0.188377 | 6.094934 | 1 | TD | 13.59843 |
| meta-analysis | 4.356249 | 0.555649 | 3.267197 | 5.445302 | 1 | TD | 100 |
| U03 | 4.461482 | 1.331367 | 1.852049 | 7.070915 | 2 | TD | 7.78695 |
| U01 | 1.443097 | 1.169078 | -0.848253 | 3.734448 | 2 | TD | 10.09895 |
| U02 | 1.529464 | 1.257414 | -0.935023 | 3.993951 | 2 | TD | 8.72984 |
| B04 | 5.870793 | 1.115440 | 3.684570 | 8.057016 | 2 | TD | 11.09356 |
| B05 | 2.429880 | 0.832626 | 0.797963 | 4.061798 | 2 | TD | 19.90964 |
| B06 | 3.929192 | 1.029053 | 1.912284 | 5.946100 | 2 | TD | 13.03429 |
| B07 | 4.526824 | 0.936627 | 2.691068 | 6.362580 | 2 | TD | 15.73366 |
| P08 | 1.890061 | 1.006939 | -0.083504 | 3.863627 | 2 | TD | 13.61308 |
| meta-analysis | 3.243405 | 0.371519 | 2.515240 | 3.971570 | 2 | TD | 100 |
| U03 | 2.778257 | 0.576077 | 1.649167 | 3.907348 | 0 | FD | 13.22894 |
| U01 | 1.395996 | 0.405790 | 0.600662 | 2.191331 | 0 | FD | 26.66137 |
| U02 | 1.507777 | 1.035791 | -0.522336 | 3.537891 | 0 | FD | 4.09205 |
| B04 | 2.169510 | 0.547698 | 1.096041 | 3.242978 | 0 | FD | 14.63537 |
| B05 | 1.354719 | 0.614957 | 0.149424 | 2.560014 | 0 | FD | 11.60902 |
| B06 | 0.394153 | 0.661359 | -0.902086 | 1.690394 | 0 | FD | 10.03717 |
| B07 | 0.208757 | 1.257328 | -2.255560 | 2.673074 | 0 | FD | 2.77708 |
| P08 | 0.897242 | 0.508795 | -0.099978 | 1.894464 | 0 | FD | 16.95896 |
| meta-analysis | 1.473733 | 0.209528 | 1.063064 | 1.884401 | 0 | FD | 100 |
| U03 | 2.120972 | 0.347484 | 1.439915 | 2.802029 | 1 | FD | 12.91694 |
| U01 | 0.757754 | 0.292544 | 0.184378 | 1.331131 | 1 | FD | 18.22411 |

| | | | | | | | |
|---|---|---|---|---|---|---|---|
| U02 | 1.305859 | 0.441987 | 0.439579 | 2.172138 | 1 | FD | 7.98382 |
| B04 | 1.800733 | 0.340406 | 1.133549 | 2.467917 | 1 | FD | 13.45969 |
| B05 | 0.687359 | 0.330652 | 0.039291 | 1.335427 | 1 | FD | 14.26545 |
| B06 | 0.308798 | 0.359360 | -0.395534 | 1.013131 | 1 | FD | 12.07731 |
| B07 | 0.181674 | 0.475525 | -0.750339 | 1.113687 | 1 | FD | 6.89734 |
| P08 | 0.860566 | 0.331702 | 0.210441 | 1.510691 | 1 | FD | 14.17531 |
| meta-analysis | 1.028557 | 0.124886 | 0.783784 | 1.273330 | 1 | FD | 100 |
| U03 | 1.758204 | 0.275799 | 1.217646 | 2.298762 | 2 | FD | 11.50275 |
| U01 | 0.465869 | 0.258250 | -0.040292 | 0.972031 | 2 | FD | 13.11919 |
| U02 | 1.189583 | 0.294767 | 0.611848 | 1.767317 | 2 | FD | 10.07001 |
| B04 | 1.544382 | 0.247923 | 1.058461 | 2.030303 | 2 | FD | 14.23493 |
| B05 | 0.400140 | 0.240097 | -0.070442 | 0.870723 | 2 | FD | 15.17796 |
| B06 | 0.384070 | 0.251713 | -0.109279 | 0.877420 | 2 | FD | 13.80946 |
| B07 | 0.159344 | 0.306976 | -0.442317 | 0.761007 | 2 | FD | 9.28497 |
| P08 | 0.808575 | 0.261443 | 0.296154 | 1.320996 | 2 | FD | 12.80068 |
| meta-analysis | 0.835063 | 0.093539 | 0.651728 | 1.018397 | 2 | FD | 100 |
| U03 | 0.314738 | 0.239154 | -0.153995 | 0.783472 | 0 | PD | 6.10241 |
| U01 | 0.315111 | 0.269988 | -0.214055 | 0.844278 | 0 | PD | 4.78816 |
| U02 | 0.569207 | 0.342518 | -0.102115 | 1.240530 | 0 | PD | 2.97502 |
| B04 | 0.403436 | 0.124217 | 0.159974 | 0.646898 | 0 | PD | 22.61997 |
| B05 | 0.158925 | 0.090812 | -0.019063 | 0.336914 | 0 | PD | 42.32222 |
| B06 | 0.460395 | 0.221982 | 0.025317 | 0.895473 | 0 | PD | 7.08305 |
| B07 | 0.413922 | 0.198960 | 0.023966 | 0.803878 | 0 | PD | 8.81705 |
| P08 | -0.163033 | 0.256812 | -0.666376 | 0.340309 | 0 | PD | 5.29207 |
| meta-analysis | 0.270224 | 0.059078 | 0.154433 | 0.386016 | 0 | PD | 100 |
| U03 | 0.037734 | 0.123886 | -0.205077 | 0.280547 | 1 | PD | 8.32787 |
| U01 | 0.713095 | 0.113593 | 0.490456 | 0.935735 | 1 | PD | 9.90540 |
| U02 | 0.513953 | 0.089485 | 0.338564 | 0.689341 | 1 | PD | 15.96145 |
| B04 | 0.388104 | 0.087382 | 0.216837 | 0.559372 | 1 | PD | 16.73892 |
| B05 | 0.082253 | 0.097049 | -0.107960 | 0.272468 | 1 | PD | 13.57032 |
| B06 | 0.674121 | 0.103127 | 0.471994 | 0.876248 | 1 | PD | 12.01788 |
| B07 | 0.432888 | 0.107759 | 0.221683 | 0.644093 | 1 | PD | 11.00698 |
| P08 | -0.119062 | 0.101236 | -0.317482 | 0.079357 | 1 | PD | 12.47114 |
| meta-analysis | 0.345753 | 0.035751 | 0.275682 | 0.415824 | 1 | PD | 100 |
| U03 | -0.046074 | 0.101350 | -0.244718 | 0.152569 | 2 | PD | 6.80732 |
| U01 | 0.759692 | 0.084543 | 0.593989 | 0.925395 | 2 | PD | 9.78286 |
| U02 | 0.354751 | 0.051880 | 0.253067 | 0.456435 | 2 | PD | 25.97909 |
| B04 | 0.376555 | 0.074171 | 0.231181 | 0.521929 | 2 | PD | 12.71019 |
| B05 | 0.065055 | 0.094957 | -0.121058 | 0.251169 | 2 | PD | 7.75476 |
| B06 | 0.651015 | 0.079457 | 0.495282 | 0.806749 | 2 | PD | 11.07552 |
| B07 | 0.359567 | 0.086213 | 0.190593 | 0.528542 | 2 | PD | 9.40771 |
| P08 | -0.066288 | 0.065133 | -0.193947 | 0.061370 | 2 | PD | 16.48249 |
| meta-analysis | 0.311254 | 0.026443 | 0.259427 | 0.363082 | 2 | PD | 100 |

*Table S2. Results from the alpha meta-analysis comparing taxonomic, functional, and phylogenetic diversity between ESBC and Control forests across study sites. The table includes effect size estimates (Difference), standard errors (SE), and confidence intervals (LCL, UCL) for each diversity order (Order q). The weight fixed column represents the relative contribution of each site to the overall meta-analysis estimate.*

| Site | Difference | SE | LCL | UCL | Order q | Diversity | weight_fixed |
|---|---|---|---|---|---|---|---|
| U01 | -2,476078 | 1,670871 | -5,750926 | 0,798769 | 0 | TD | 14,850500 |
| U02 | -0,288350 | 1,775658 | -3,768576 | 3,191876 | 0 | TD | 13,149467 |
| U03 | -0,354091 | 2,143503 | -4,555279 | 3,847097 | 0 | TD | 9,023581 |
| B04 | 7,194106 | 1,583153 | 4,091183 | 10,297028 | 0 | TD | 16,541741 |
| B05 | 7,441869 | 1,872678 | 3,771488 | 11,112250 | 0 | TD | 11,822268 |
| B06 | 1,868642 | 3,321800 | -4,641966 | 8,379250 | 0 | TD | 3,757336 |
| B07 | 5,891358 | 1,602881 | 2,749769 | 9,032947 | 0 | TD | 16,137063 |
| P08 | -0,241701 | 1,678373 | -3,531251 | 3,047849 | 0 | TD | 14,718045 |
| meta_analysis | 2,617579 | 0,643893 | 1,355573 | 3,879586 | 0 | TD | 100 |
| U01 | -1,575907 | 1,419050 | -4,357194 | 1,205380 | 1 | TD | 9,213912 |
| U02 | -0,983902 | 1,269176 | -3,471441 | 1,503636 | 1 | TD | 11,518502 |
| U03 | 1,469851 | 1,518570 | -1,506492 | 4,446193 | 1 | TD | 8,045813 |
| B04 | 6,418163 | 1,193943 | 4,078078 | 8,758248 | 1 | TD | 13,015850 |
| B05 | 4,932766 | 1,323452 | 2,338848 | 7,526685 | 1 | TD | 10,593098 |
| B06 | 2,789893 | 1,409424 | 0,027473 | 5,552313 | 1 | TD | 9,340202 |
| B07 | 4,992741 | 0,885022 | 3,258130 | 6,727352 | 1 | TD | 23,688170 |
| P08 | 0,091377 | 1,127911 | -2,119287 | 2,302041 | 1 | TD | 14,584453 |
| meta_analysis | 2,674237 | 0,430745 | 1,829993 | 3,518480 | 1 | TD | 100 |
| U01 | -0,223031 | 0,927732 | -2,041353 | 1,595291 | 2 | TD | 10,213143 |
| U02 | -0,990836 | 0,924212 | -2,802258 | 0,820585 | 2 | TD | 10,291104 |
| U03 | 2,734705 | 0,805849 | 1,155270 | 4,314141 | 2 | TD | 13,536212 |
| B04 | 5,972022 | 0,995464 | 4,020950 | 7,923095 | 2 | TD | 8,870622 |
| B05 | 3,092058 | 0,954092 | 1,222072 | 4,962045 | 2 | TD | 9,656597 |
| B06 | 3,485801 | 0,834141 | 1,850915 | 5,120687 | 2 | TD | 12,633569 |
| B07 | 4,249439 | 0,626825 | 3,020885 | 5,477994 | 2 | TD | 22,372371 |
| P08 | 0,387232 | 0,841066 | -1,261227 | 2,035691 | 2 | TD | 12,426382 |
| meta_analysis | 2,512973 | 0,296485 | 1,931873 | 3,094072 | 2 | TD | 100 |
| U01 | -0,404328 | 1,443703 | -3,233933 | 2,425277 | 0 | FD | 12,152992 |
| U02 | 0,965197 | 1,941849 | -2,840757 | 4,771152 | 0 | FD | 6,717500 |
| U03 | 0,780463 | 1,606479 | -2,368178 | 3,929104 | 0 | FD | 9,814961 |
| B04 | 6,026689 | 1,382653 | 3,316740 | 8,736638 | 0 | FD | 13,249896 |
| B05 | 3,848096 | 1,451910 | 1,002406 | 6,693787 | 0 | FD | 12,015987 |
| B06 | 3,229344 | 1,503956 | 0,281644 | 6,177043 | 0 | FD | 11,198722 |
| B07 | 5,871560 | 1,230267 | 3,460280 | 8,282840 | 0 | FD | 16,735538 |
| P08 | 1,823230 | 1,182517 | -0,494460 | 4,140921 | 0 | FD | 18,114403 |
| meta_analysis | 3,027768 | 0,503291 | 2,041335 | 4,014200 | 0 | FD | 100 |
| U01 | -0,182539 | 1,431377 | -2,987987 | 2,622909 | 1 | FD | 8,498795 |
| U02 | 0,288013 | 1,485551 | -2,623613 | 3,199638 | 1 | FD | 7,890251 |
| U03 | 1,210479 | 1,324146 | -1,384799 | 3,805756 | 1 | FD | 9,931028 |
| B04 | 5,335679 | 1,089135 | 3,201013 | 7,470345 | 1 | FD | 14,679187 |
| B05 | 2,806447 | 1,132582 | 0,586627 | 5,026266 | 1 | FD | 13,574587 |

| | | | | | | | |
|---|---|---|---|---|---|---|---|
| B06 | 3,130119 | 1,171695 | 0,833639 | 5,426598 | 1 | FD | 12,683428 |
| B07 | 5,374841 | 1,040996 | 3,334526 | 7,415155 | 1 | FD | 16,068210 |
| P08 | 1,205405 | 1,021895 | -0,797472 | 3,208282 | 1 | FD | 16,674513 |
| meta_analysis | 2,753265 | 0,417285 | 1,935401 | 3,571128 | 1 | FD | 100 |
| U01 | 0,210624 | 1,229546 | -2,199242 | 2,620490 | 2 | FD | 10,106316 |
| U02 | 0,432647 | 1,059590 | -1,644111 | 2,509405 | 2 | FD | 13,608389 |
| U03 | 1,519218 | 1,166148 | -0,766390 | 3,804825 | 2 | FD | 11,235061 |
| B04 | 4,967287 | 1,298613 | 2,422052 | 7,512522 | 2 | FD | 9,059889 |
| B05 | 2,351058 | 1,070379 | 0,253155 | 4,448962 | 2 | FD | 13,335444 |
| B06 | 3,146753 | 1,339237 | 0,521896 | 5,771609 | 2 | FD | 8,518585 |
| B07 | 5,120283 | 0,963051 | 3,232738 | 7,007827 | 2 | FD | 16,473440 |
| P08 | 0,934388 | 0,930059 | -0,888494 | 2,757271 | 2 | FD | 17,662876 |
| meta_analysis | 2,290988 | 0,390878 | 1,524881 | 3,057095 | 2 | FD | 100 |
| U01 | -0,142130 | 0,344530 | -0,817395 | 0,533136 | 0 | PD | 9,489653 |
| U02 | 0,301374 | 0,267781 | -0,223466 | 0,826215 | 0 | PD | 15,708869 |
| U03 | -0,132627 | 0,335170 | -0,789549 | 0,524295 | 0 | PD | 10,027021 |
| B04 | 0,021514 | 0,264125 | -0,496161 | 0,539190 | 0 | PD | 16,146729 |
| B05 | 0,021276 | 0,264183 | -0,496514 | 0,539066 | 0 | PD | 16,139584 |
| B06 | -0,239577 | 0,336935 | -0,899957 | 0,420804 | 0 | PD | 9,922269 |
| B07 | 0,452161 | 0,280021 | -0,096671 | 1,000993 | 0 | PD | 14,365516 |
| P08 | -0,426023 | 0,370626 | -1,152436 | 0,300389 | 0 | PD | 8,200359 |
| meta_analysis | 0,033712 | 0,106133 | -0,174305 | 0,241730 | 0 | PD | 100 |
| U01 | 0,420916 | 0,164954 | 0,097611 | 0,744220 | 1 | PD | 11,169556 |
| U02 | 0,370887 | 0,146458 | 0,083833 | 0,657940 | 1 | PD | 14,168851 |
| U03 | -0,166878 | 0,175465 | -0,510782 | 0,177027 | 1 | PD | 9,871533 |
| B04 | 0,266041 | 0,176956 | -0,080786 | 0,612868 | 1 | PD | 9,705847 |
| B05 | 0,044294 | 0,141338 | -0,232724 | 0,321312 | 1 | PD | 15,213987 |
| B06 | 0,566071 | 0,139460 | 0,292734 | 0,839407 | 1 | PD | 15,626579 |
| B07 | 0,401128 | 0,146071 | 0,114834 | 0,687422 | 1 | PD | 14,244105 |
| P08 | -0,210743 | 0,174338 | -0,552438 | 0,130953 | 1 | PD | 9,999542 |
| meta_analysis | 0,240173 | 0,055129 | 0,132122 | 0,348224 | 1 | PD | 100 |
| U01 | 0,564554 | 0,120206 | 0,328954 | 0,800153 | 2 | PD | 13,421635 |
| U02 | 0,361922 | 0,115368 | 0,135805 | 0,588038 | 2 | PD | 14,571064 |
| U03 | -0,156890 | 0,158953 | -0,468431 | 0,154651 | 2 | PD | 7,675807 |
| B04 | 0,377954 | 0,123993 | 0,134932 | 0,620976 | 2 | PD | 12,614301 |
| B05 | 0,048407 | 0,117475 | -0,181839 | 0,278654 | 2 | PD | 14,053015 |
| B06 | 0,653142 | 0,142841 | 0,373178 | 0,933106 | 2 | PD | 9,504958 |
| B07 | 0,339278 | 0,120580 | 0,102946 | 0,575610 | 2 | PD | 13,338627 |
| P08 | -0,130071 | 0,114392 | -0,354276 | 0,094133 | 2 | PD | 14,820591 |
| meta_analysis | 0,259003 | 0,044038 | 0,172690 | 0,345316 | 2 | PD | 100 |

*Table S3. Results from the beta 1-S meta-analysis comparing taxonomic, functional, and phylogenetic diversity between ESBC and Control forests across study sites. The table includes effect size estimates (Difference), standard errors (SE), and confidence intervals (LCL, UCL) for each diversity order (Order q). The weight fixed column represents the relative contribution of each site to the overall meta-analysis estimate.*

| Site | Difference | SE | LCL | UCL | Order q | Diversity | weight_fixed |
|---|---|---|---|---|---|---|---|
| U01 | 0,512062 | 0,037777 | 0,438020 | 0,586103 | 0 | TD | 16,514059 |
| U02 | -0,091057 | 0,025622 | -0,141275 | -0,040839 | 0 | TD | 35,898736 |
| U03 | 0,100715 | 0,042740 | 0,016947 | 0,184484 | 0 | TD | 12,901634 |
| B04 | 0,123177 | 0,059043 | 0,007455 | 0,238898 | 0 | TD | 6,760438 |
| B05 | -0,046046 | 0,058835 | -0,161360 | 0,069267 | 0 | TD | 6,808372 |
| B06 | -0,004378 | 0,050143 | -0,102656 | 0,093900 | 0 | TD | 9,373281 |
| B07 | 0,198379 | 0,060040 | 0,080703 | 0,316055 | 0 | TD | 6,537711 |
| P08 | 0,535956 | 0,067284 | 0,404082 | 0,667830 | 0 | TD | 5,205767 |
| meta_analysis | 0,110520 | 0,015352 | 0,080431 | 0,140608 | 0 | TD | 100 |
| U01 | 0,381163 | 0,029002 | 0,324321 | 0,438005 | 1 | TD | 19,659855 |
| U02 | 0,002127 | 0,029287 | -0,055274 | 0,059528 | 1 | TD | 19,279010 |
| U03 | 0,011600 | 0,033437 | -0,053935 | 0,077135 | 1 | TD | 14,790071 |
| B04 | 0,091292 | 0,039789 | 0,013306 | 0,169278 | 1 | TD | 10,444466 |
| B05 | 0,000432 | 0,036280 | -0,070676 | 0,071539 | 1 | TD | 12,562844 |
| B06 | 0,010906 | 0,041071 | -0,069592 | 0,091405 | 1 | TD | 9,802604 |
| B07 | 0,110922 | 0,049182 | 0,014527 | 0,207317 | 1 | TD | 6,836103 |
| P08 | 0,418351 | 0,049959 | 0,320433 | 0,516269 | 1 | TD | 6,625048 |
| meta_analysis | 0,123019 | 0,012859 | 0,097815 | 0,148222 | 1 | TD | 100 |
| U01 | 0,193202 | 0,021285 | 0,151484 | 0,234919 | 2 | TD | 22,562734 |
| U02 | 0,105221 | 0,020328 | 0,065380 | 0,145063 | 2 | TD | 24,737794 |
| U03 | -0,084265 | 0,033581 | -0,150081 | -0,018448 | 2 | TD | 9,064716 |
| B04 | 0,060131 | 0,036086 | -0,010596 | 0,130857 | 2 | TD | 7,849881 |
| B05 | 0,043278 | 0,031164 | -0,017802 | 0,104358 | 2 | TD | 10,525292 |
| B06 | -0,012526 | 0,027620 | -0,066660 | 0,041607 | 2 | TD | 13,399648 |
| B07 | 0,049215 | 0,037070 | -0,023441 | 0,121871 | 2 | TD | 7,438449 |
| P08 | 0,306337 | 0,048082 | 0,212098 | 0,400576 | 2 | TD | 4,421486 |
| meta_analysis | 0,086785 | 0,010110 | 0,066969 | 0,106601 | 2 | TD | 100 |
| U01 | 0,309881 | 0,059220 | 0,193811 | 0,425950 | 0 | FD | 13,996154 |
| U02 | -0,029067 | 0,082150 | -0,190079 | 0,131945 | 0 | FD | 7,273250 |
| U03 | 0,035941 | 0,070332 | -0,101906 | 0,173789 | 0 | FD | 9,923062 |
| B04 | 0,032843 | 0,046287 | -0,057878 | 0,123564 | 0 | FD | 22,910351 |
| B05 | -0,044885 | 0,044262 | -0,131638 | 0,041868 | 0 | FD | 25,053976 |
| B06 | 0,006506 | 0,085933 | -0,161919 | 0,174931 | 0 | FD | 6,647067 |
| B07 | 0,056264 | 0,073009 | -0,086831 | 0,199359 | 0 | FD | 9,208603 |
| P08 | 0,205139 | 0,099204 | 0,010702 | 0,399576 | 0 | FD | 4,987536 |
| meta_analysis | 0,056948 | 0,022155 | 0,013524 | 0,100371 | 0 | FD | 100 |
| U01 | 0,190167 | 0,037916 | 0,115854 | 0,264480 | 1 | FD | 11,083379 |
| U02 | 0,093492 | 0,039767 | 0,015549 | 0,171434 | 1 | FD | 10,075302 |
| U03 | -0,014413 | 0,049169 | -0,110783 | 0,081957 | 1 | FD | 6,590489 |

| | | | | | | | |
|---|---|---|---|---|---|---|---|
| B04 | 0,038231 | 0,023688 | -0,008196 | 0,084658 | 1 | FD | 28,396655 |
| B05 | -0,001031 | 0,028549 | -0,056985 | 0,054923 | 1 | FD | 19,549606 |
| B06 | 0,020254 | 0,046892 | -0,071653 | 0,112160 | 1 | FD | 7,246247 |
| B07 | -0,002388 | 0,035170 | -0,071320 | 0,066545 | 1 | FD | 12,881186 |
| P08 | 0,196159 | 0,061761 | 0,075110 | 0,317208 | 1 | FD | 4,177136 |
| meta_analysis | 0,049555 | 0,012623 | 0,024815 | 0,074295 | 1 | FD | 100 |
| U01 | 0,080406 | 0,041519 | -0,000968 | 0,161781 | 2 | FD | 5,378677 |
| U02 | 0,125425 | 0,035628 | 0,055595 | 0,195254 | 2 | FD | 7,304248 |
| U03 | -0,073733 | 0,049837 | -0,171411 | 0,023946 | 2 | FD | 3,732996 |
| B04 | 0,045838 | 0,016064 | 0,014354 | 0,077323 | 2 | FD | 35,930391 |
| B05 | 0,028256 | 0,018190 | -0,007396 | 0,063908 | 2 | FD | 28,021485 |
| B06 | -0,000438 | 0,028291 | -0,055888 | 0,055013 | 2 | FD | 11,583708 |
| B07 | -0,036165 | 0,040157 | -0,114870 | 0,042541 | 2 | FD | 5,749704 |
| P08 | 0,173684 | 0,063508 | 0,049210 | 0,298158 | 2 | FD | 2,298790 |
| meta_analysis | 0,036984 | 0,009629 | 0,018111 | 0,055856 | 2 | FD | 100 |
| U01 | 0,255271 | 0,088461 | 0,081890 | 0,428651 | 0 | PD | 19,147058 |
| U02 | -0,003988 | 0,095184 | -0,190544 | 0,182568 | 0 | PD | 16,538010 |
| U03 | -0,001417 | 0,093162 | -0,184010 | 0,181176 | 0 | PD | 17,263723 |
| B04 | 0,310553 | 0,113065 | 0,088950 | 0,532156 | 0 | PD | 11,720684 |
| B05 | -0,031703 | 0,098309 | -0,224385 | 0,160979 | 0 | PD | 15,503185 |
| B06 | 0,294786 | 0,129116 | 0,041723 | 0,547849 | 0 | PD | 8,987651 |
| B07 | -0,110785 | 0,228904 | -0,559428 | 0,337858 | 0 | PD | 2,859573 |
| P08 | 0,252903 | 0,137025 | -0,015661 | 0,521466 | 0 | PD | 7,980116 |
| meta_analysis | 0,122965 | 0,038708 | 0,047098 | 0,198832 | 0 | PD | 100 |
| U01 | -0,017254 | 0,053397 | -0,121910 | 0,087403 | 1 | PD | 23,803175 |
| U02 | -0,018575 | 0,061555 | -0,139221 | 0,102072 | 1 | PD | 17,911809 |
| U03 | 0,005116 | 0,064622 | -0,121542 | 0,131774 | 1 | PD | 16,251923 |
| B04 | 0,126170 | 0,103905 | -0,077480 | 0,329821 | 1 | PD | 6,286323 |
| B05 | -0,023061 | 0,062708 | -0,145967 | 0,099844 | 1 | PD | 17,259381 |
| B06 | -0,107359 | 0,081548 | -0,267191 | 0,052473 | 1 | PD | 10,205640 |
| B07 | -0,131381 | 0,134418 | -0,394835 | 0,132073 | 1 | PD | 3,756274 |
| P08 | 0,012298 | 0,122463 | -0,227725 | 0,252320 | 1 | PD | 4,525474 |
| meta_analysis | -0,017986 | 0,026052 | -0,069047 | 0,033074 | 1 | PD | 100 |
| U01 | -0,089528 | 0,051021 | -0,189526 | 0,010471 | 2 | PD | 21,957676 |
| U02 | 0,014027 | 0,065624 | -0,114594 | 0,142649 | 2 | PD | 13,272366 |
| U03 | 0,009313 | 0,050750 | -0,090155 | 0,108781 | 2 | PD | 22,192489 |
| B04 | 0,017642 | 0,076294 | -0,131892 | 0,167175 | 2 | PD | 9,819723 |
| B05 | -0,013629 | 0,069151 | -0,149163 | 0,121905 | 2 | PD | 11,953126 |
| B06 | -0,196457 | 0,063602 | -0,321115 | -0,071799 | 2 | PD | 14,129685 |
| B07 | -0,099148 | 0,111475 | -0,317635 | 0,119338 | 2 | PD | 4,599660 |
| P08 | -0,072961 | 0,165959 | -0,398236 | 0,252313 | 2 | PD | 2,075275 |
| meta_analysis | -0,049460 | 0,023908 | -0,096318 | -0,002601 | 2 | PD | 100 |

*Table S4. Functional traits and trait states of hoverflies sampled.*

| Functional traits | Functional trait states | Description |
|---|---|---|
| Larval microhabitat | - Trees<br>- Upward climbing lianas<br>- Herb layer<br>- Timber<br>- Dung<br>- Litter<br>- Stones<br>- Nests of social insects<br>- Root zone<br>- On water plants<br>- Submerged sediment | Defines microhabitat in which larvae develop. 11 trait states, not mutually exclusive. |
| Larval food type | - Saprophagous<br>- Saproxylic<br>- Zoophagous<br>- Phytophagous related to roots<br>- Phytophagous related to bulbs | Feeding preference of larvae. 5 trait states. |
| Duration of larval development | - Less than 2 months<br>- 2-6 months<br>- 7-12 months<br>- More than a year | 4 traits states, not mutually exclusive. |
| Number of generations during the year | - Less than one<br>- One<br>- Two<br>- More than two | 4 trait states, not mutually exclusive. |
| Inundation tolerance of larvae | - Non tolerant<br>- Tolerant with long breathing tube<br>- Tolerant with short breathing tube | Indicates the inundation tolerance of larvae. 4 trait states. If it's tolerant it is specified if the breathing tube is short, medium or long. |

| | | |
|---|---|---|
| | - Tolerant with breathing tube of medium length | |
| Fight period of adults | - Early spring<br>- Spring<br>- Early summer<br>- Summer<br>- Autumn | 5 trait states, not mutually exclusive. |
| Distribution | - North distribution<br>- South distribution<br>- Widely distributed | Indicates if the species has predominantly northern, southern distribution in Europe, or if it's widely distributed. |
| Body size of adults | - Small (less than 7 mm)<br>- Small or medium<br>- Medium (7-12 mm)<br>- Medium or robust<br>- Robust (larger than 12 mm) | Species were categorized into 3 groups based on their size. 5 trait states are present, as the species could cover the range of more than one category. |
| Flying ability | - Bad<br>- Good<br>- Very good | 3 trait states: bad (for the species flying clumsily and over very short distances), good (flying freely across the area), very good (for the species with migrating potential) |
| Adult microhabitat | - temperate deciduous forests,<br>- Mediterranean forests and bushes<br>- Coniferous forests<br>- Alpine habitats<br>- Wetland habitats<br>- Steppe habitats<br>- Not specialized | 7 trait states, not mutually exclusive |
| Human impact tolerance | - Low<br>- Medium<br>- High | Defines the sensitivity of adults to human impact tollerance. 4 trait states: low (natural or only slightly |

| | | |
|---|---|---|
| | - Very high | changed habitat), medium (low grazing, ecological friendly forestry), high (over-grazing, forest cutting, replacement of initial forest, agriculture, villages), very high (destruction of habitat, intense agriculture as monoculture without borders, cities) |
